# Two Interaction Surfaces between XPA and RPA Organize the Preincision Complex in Nucleotide Excision Repair

**DOI:** 10.1101/2022.03.01.482439

**Authors:** Mihyun Kim, Hyun Suk Kim, Areetha D’Souza, Kaitlyn Gallagher, Eunwoo Jeong, Agnieszka Topolska-Wos, Kateryna Ogorodnik Le Meur, Chi-Lin Tsai, Miaw-Sheue Tsai, Minyong Kee, John A. Tainer, Jung-Eun Yeo, Walter J. Chazin, Orlando D. Schärer

## Abstract

The XPA and RPA proteins fulfill essential roles in the assembly of the preincision complex in the nucleotide excision repair pathway. We have previously characterized the two interaction surfaces between XPA and RPA, with the RPA32 and RPA70AB subunits. Here we show that the mutations in the two individual interaction surfaces reduce NER activity in biochemical and cellular systems, and that combining mutations in two domains leads to an additive inhibition of NER, suggesting that they fulfill distinct roles. Our data suggest that the interaction between XPA and RPA32 is important for the initial association of XPA with NER complexes, while the interaction between XPA and RPA70 is needed for structural organization of the complex to license the dual incision reaction. SAXS analysis of complexes of XPA and RPA bound to ss/dsDNA junction substrates reveals the architecture of XPA and RPA in the preincision complex and shows that the two interaction domains between RPA and XPA are located at opposite sides of the two molecules. We propose a structure for the overall NER preincision complex that shows that the two strands of the NER bubble assume a U-shape with the two ss/dsDNA junctions localized in close proximity, with the interaction between XPA and RPA70 as one of the key organizing elements.

## Introduction

Nucleotide excision repair (NER) is the primary repair pathway for the removal of bulky DNA adducts induced by UV irradiation, environmental mutagens, and anticancer agents from our genomes ^1^. People with mutations in NER genes suffer from xeroderma pigmentosum (XP), a disease characterized by extreme sensitivity to UV irradiation and a highly increased incidence of skin cancer ^2^. NER is multistep pathway involving more than 30 proteins and is initiated by XPC-RAD23B, which recognizes aberrations in duplex DNA ^3^. For some lesions, in particular cyclopyrimidine dimers (CPDs) in chromatin, this recognition additionally requires UV-DDB and its associated ubiquitin ligase complex ^4^. XPC-RAD23B recruits the 10 subunit complex TFIIH to lesions ^5^, which, using the XPB translocase and XPD helicase, opens the DNA and verifies the lesion ^6,7^. XPA and RPA are then recruited to the opened DNA ^5,8^, stabilize the pre-incision complex and help to position the ERCC1-XPF and XPG endonucleases ^9^. ERCC1-XPF first incises the DNA 5’ to the lesion, then replication proteins initiate gap filling, and following 3’ incision by XPG, gap filling is completed and resulting nick sealed ^10-12^. Progression through the NER pathway is driven by multiple protein-protein and protein-DNA interactions among the factors and substrates ^13,14^.

The focus of this work is the critical role of XPA and RPA in organizing the pre-incision complex and licensing the dual incision reaction. XPA is a small protein consisting of only 273 residues that acts as scaffold by interacting with several NER proteins and the DNA substrate ^15^. XPA has a central globular core and disordered N- and C-termini. XPA interacts with the opened NER bubble structure at the 5’ ssDNA–dsDNA junction via its central domain and binding to the adjacent duplex with a long C-terminal α helix extending from the central domain ^16-19^. The binding of XPA to DNA stimulates interactions with NER proteins, including RPA ^20-24^, and ERCC1-XPF ^25,26^.

RPA, the ubiquitous eukaryotic ssDNA binding protein, is a heterotrimer consisting of the RPA70, RPA32, and RPA14 subunits ^27^. The RPA ssDNA binding apparatus consists of 4 OB-fold domains (RPA70A, RPA70B, RPA70C and RPA32D) and two recruitment domains on long flexible linkers: OB-fold domain RPA70N and winged helix domain RPA32C. XPA binds to both RPA32C and RPA70AB ^20,24,28^. Using pull down assays, it was reported that residues 4-29 of XPA bind to RPA32 ^21^. However, NMR analysis revealed the RPA32C-binding motif as XPA_29-46_, a region that also binds other DNA repair proteins UNG2 and RAD52 ^20^. XPA has a second weaker interaction with RPA70AB through its DNA binding domain (DBD). This interaction was originally attributed to XPA residues 153-176 ^23,29^, and certain mutations in that region result in reduced NER activity. However, NMR chemical shift perturbation analysis mapped RPA70AB interaction to XPA residues 98-129 ^22^. Recently, we reported a structural model that shows that residues 101-114 in and around the XPA-DBD Zn binding motif interact with RPA70AB. Mutations in several of these residues resulted in reduction in binding affinity and *in vitro* NER activity ^24^. Thus, the XPA zinc binding motif interacts with both RPA70AB and the ssDNA-dsDNA junction in the NER bubble substrate.

Here, we report investigations of how the two contacts between XPA and RPA contribute to the NER reaction using a structural approach and mutations in XPA that disrupt the binding interface with both RPA32C and RPA70AB. Biochemical and cellular assays show synergistic contributions to the physical interaction between RPA and XPA and to overall NER activity. Our results support a model in which the two contacts between XPA and RPA can be engaged simultaneously and are critical to the progression of the NER pathway.

## Materials and Methods

### Plasmids and antibodies

HA tagged XPA cDNA was cloned into a pWPXL vector. Mutated pWPXL-XPA and pBG100-XPA expression vectors were generated by site-directed mutagenesis from pWPXL-XPA WT and pBG100-XPA WT using the KOD-mutagenesis kit (Toyobo, TYB-SMK-101). The following primers were used for inverse PCR mutagenesis (altered codons underlined):

XPA 70-M1K_F: 5’-<u>AAA</u>GAATGTGGGAAAGAATTTATGG-3’

XPA 70-M1K_R: 5’-GCATATTACATAATCAAATTCCATAACAGGTCC-3’

XPA 70-M1A_F: 5’-GAA<u>GCT</u>ATGGATTCTTATCTTATGAACCAC-3’

XPA 70-M1A_R: 5’-TTTCCCACATTCTTCGCATATTACA-3’

XPA 32-E1_F (R34E): 5’-<u>GAA</u>GCACTGATGCTGCGCCAGGCC-3’

XPA 32-E1_R (R34E): 5’-CTGCCGCTTCCGCTCGATACTCG-3’

(XPA 70-M2_F primer is same with XPA 70-M1A_F primer.)

XPA 70-M2_R: 5’-TTTCCCACATTCTT<u>TGC</u>ATATTACA-3’

XPA 70-M3_F (D101N/E106K/F112A): 5’-<u>AAT</u>TATGTAATATGC<u>AAA</u>GAATGTGGGAA-3’

XPA 70-M3_R (D101N/E106K/F112A): 5’-AAATTCCATAACAGGTCCTGGTT-3’

XPA 70-M4_F (D101/E106K/K110E/F112A): 5’-GG<u>GAA</u>GAA<u>GCT</u>ATGGATTCTTATC-3’

XPA 70-M4_R (D101/E106K/K110E/F112A): 5’-CACATTC<u>TTT</u>GCATATTACATA<u>ATT</u>AAA-3’

XPA 70-M6_F (D101N/E106K/K110E/E111K/F112A/D114N): 5’-<u>AAAGCT</u>ATG<u>AAT</u>TCTTATCTTATGAACC-3’

XPA 70-M6_R (D101N/E106K/K110E/E111K/F112A/D114N): 5’-<u>TTC</u>CCCACATTC<u>TTT</u>GCATATTAC-3’.

XPA 32-E3_F (R34E/R39E/R42E): 5’-GCC<u>GAA</u>CTGGCTGCCCGGCCCTA-3’

XPA 32-E3_R (R34E/R39E/R42E): 5’-CTG<u>TTC</u>CAGCATCAGTGCTTCCT-3’

XPA 32-W4_F (R30E/K31E/R32E/R34E): 5’-<u>GAAGAA</u>CAGGAAGCACTGATGCTGC-3’

XPA 32-E4_R (R30E/K31E/R32E/R34E): 5’-<u>TTC</u>CTCGATACTCGCCCGCACCGA-3’

(XPA 32-E6_F (R30E/K31E/R32E/R34E/R39E/R42E) primer is same with XPA 32-E3_F primer. XPA 32-E6_R (R30E/K31E/R32E/R34E/R39E/R42E) primer is same with XPA 32-E3_R primer).

The following antibodies were used for Western blot: anti-XPA (Santacruz, sc-853), anti-HA (Abcam, ab9110), Ku80 (Cell signaling, 2753s), and Goat anti-rabbit IgG (Enzo, ADI-SAB-300). The following antibodies were used for local UV irradiation assay and slot-blot assays: Anti-(6-4) PP (Cosmo bio, CAC-NM-DND-002), anti-CPD (Cosmo bio, CAC-NM-DND-001), and Cy™3 AffiniPure Goat Anti-Mouse IgG (Jackson immunoresearch, 115-165-146).

### XPA Protein expression and purification

Wild-type or mutant full-length XPA with an N-terminal His_6_ tag was expressed in *Escherichia coli* Rosetta pLysS cells. Cells were grown in TB medium containing 50 µg/ml kanamycin to OD_600_ = 0.6 at 37 °C, then OD_600_ = 1.2 at 18 °C, followed by induction with 0.5 mM isopropyl β-D-thiogalactoside at 18 °C overnight. Cells were collected by centrifugation at 6,500 rpm for 20 min and all subsequent purification steps were performed at 4 °C. The cell pellet from 1 L of culture was resuspended in 20 mL of lysis buffer (100 mM Tris-HCl at pH 8.0, 500 mM NaCl, 20 mM imidazole, 5 mM 2-mercaptoethanol, 10 µM ZnCl_2_, 200 µg/ml lysozyme, 0.5 mM PMSF, 1 mM benzamidine, and 10% glycerol, and protease inhibitor (Roche)). The suspended pellets were Dounce homogenized 20 times and sonicated with an amplitude of 60 (5 sec on/10 sec off) for 2 min. The lysate was clarified by centrifugation at 20,000 rpm for 40 min and filtration of the supernatant through a 0.45 mm syringe filter. The supernatant was incubated for 90 min at 4 °C with 10 mL of Ni-resin pre-equilibrated with Ni-loading buffer (100 mM Tris-HCl at pH 8.0, 500 mM NaCl, 20 mM imidazole, 5 mM 2-mercaptoethanol, 10 µM ZnCl_2_, and 10% glycerol). The resin was washed twice with Ni-wash buffer (100 mM Tris-HCl at pH 8.0, 500 mM NaCl, 20 mM imidazole, 5 mM 2-mercaptoethanol, 10 µM ZnCl_2_, and 10% glycerol) and eluted with Ni-elution buffer (100 mM Tris-HCl at pH 8.0, 500 mM NaCl, 400 mM imidazole, 5 mM 2-mercaptoethanol, 10 µM ZnCl_2_, and 10% glycerol). The eluent was incubated with H3C protease overnight at 4°C and dialyzed in buffer containing 20 mM Tris-HCl at pH 8.0, 300 mM NaCl, 5 mM 2-mercaptoethanol, 10 µM ZnCl_2_, and 10% glycerol. The dialyzed sample was diluted to a concentration of 150 mM NaCl using a buffer of 20 mM Tris-HCl at pH 8.0, 5 mM 2-mercaptoethanol, 10 µM ZnCl_2_, and 10% glycerol, then applied to a 5 mL Hi-trap heparin column equilibrated with Hep-buffer (20 mM Tris-HCl at pH 8.0, 50 mM NaCl, 5 mM 2-mercaptoethanol, 10 µM ZnCl_2_, and 10% glycerol) and eluted using a linear gradient with Hep/NaCl-buffer (20 mM Tris-HCl at pH 8.0, 1 M NaCl, 5 mM 2-mercaptoethanol, 10 µM ZnCl_2_, and 10% glycerol) over 5 column volumes. The XPA protein eluted at approximately 400 mM NaCl. The proteins were further purified on a HiLoad 16/600 Superdex 75pg column with buffer (50 mM Tris-HCl at pH 8.0, 150 mM NaCl, 1 mM dithiothreitol, 10 µM ZnCl_2_, and 10% glycerol), where they eluted at 50-55 ml. The proteins were obtained at a concentration of ∼1 mg/mL, yielding a total of 2-3 mg per liter of cell culture.

### Electrophoretic mobility shift assay (EMSA)

EMSA was conducted using three-way junction substrate as described in our previous paper ^24^. The annealed three-way junction oligonucleotides (100nM) with fluorescein-label were incubated with wild-type or mutant XPA (0, 20, 40, 80 nM) in a 10 µL mixture containing 25 mM Tris-HCl (pH 8.0), 1 mM DTT, 0.1 mg/ml BSA, 5% glycerol, and 1 mM EDTA at 25 °C for 30 min. The reaction mixture was loaded onto 8% native polyacrylamide gel and run at 4 °C for 2 hr at 20 mA with 0.5% TBE buffer. Gels were scanned using an Amersham Typhoon RGB imager. The band intensity was measured with ImageQuant TL program and the fraction of bound XPA-bound protein quantified based on the intensity of the free DNA band.

### Isothermal Titration Calorimetry (ITC)

Protein samples were dialyzed in 20 mM Tris (pH 8.0), 150 mM NaCl, 3% glycerol, and 0.5 mM TCEP. Experiments were performed at 25 °C with 125 rpm stirring using an Affinity ITC instrument (TA Instruments). Titrations were performed by titrating an initial injection of 1 µL of 1.1 × 10^−4^ M XPA into the sample cell containing 0.20 × 10^−4^ M RPA, followed by an additional 47 injections of 3 µL each. The injections were spaced over 200-250 s intervals. Data were analyzed using NanoAnalyze software provided by TA Instruments. The thermodynamic parameters and binding affinities (K_d_) were calculated using the average of three titrations fit to an independent binding model.

The XPA E106K, F112A mutant is unstable at high concentrations and forms a precipitate during titrations. The precipitate caused mechanical issues during one of the replicates, and the last 10 injections were removed for analysis. To validate the K_d_ and stoichiometry calculations of the three forward titrations for the 70AB-E106K, F112A mutant, we performed a reverse titration. The titration was conducted by injecting 1 µL of 90 × 10^−6^ M RPA into the sample cell containing 16 × 10^−6^ M XPA 70AB-E106K, F112A followed by an additional 47 injections of 3 µL each. Thermodynamic parameters similar to the forward titration were obtained.

### Microscale Thermophoresis (MST)

DNA-binding affinity was measured using a Monolith NT.115 (NanoTemper Technologies). The XPA samples were buffer exchanged into buffer containing 50 mM Tris-HCl (pH 7.8), 150 mM NaCl, 10 mM MgCl_2_, 0.05% Tween-20, and 1 mM DTT and diluted into 16 concentrations ranging from 0.1 to 50 × 10^−6^ M. A 5’-6-FAM labelled 4-nucleotide hairpin substrate (5’–TTTTGCGGCCGCTTTTGCGGCCGC–3’) was added to the protein at a final concentration of 40 × 10^−9^ M. All measurements were carried out at room temperature in standard capillaries (Nanotemper Technologies, Cat# MO-K022) using 20% excitation power and 20% MST power. K_d_ values were determined by fitting the data to the One Site – Total Binding model in GraphPad Prism version 8. Outlier data points were identified using a box plot analysis and removed if outside the whiskers.

### Small-angle X-ray scattering (SAXS)

The SAXS profiles were collected in SEC-SAXS mode at the ALS beamline 12.3.1 LBNL Berkeley, California ^30^. The X-ray wavelength λ was 1.03 Å and the sample-to-detector distance was set to 1.5 m resulting in scattering vectors, q, ranging from 0.01 Å^-1^ to 0.5 Å^-1^. The scattering vector is defined as q = 4π sinθ/λ, where 2θ is the scattering angle. All experiments were performed at 20 °C ^31^ and data were processed as described ^32^. Briefly, the flow through SAXS cell was directly coupled with an online Agilent 1260 Infinity HPLC system using a Shodex KW803 column (Shodex™). The column was equilibrated with running buffer (20 mM Tris pH 8.0, 150 mM NaCl, 2% glycerol, 1 mM DTT) with a flow rate of 0.5 mL/min. A 50 µL sample of the complex (prepared in 1:1:1 stoichiometry with a final concentration of 25μM) was run through the SEC and 3.0 second X-ray exposures were collected continuously during a ∼35 min elution. The SAXS frames recorded prior to the protein elution peak were used as buffer blanks to subtract from all other frames. The subtracted frames were examined by radius of gyration (GR_g_) and scattering intensity at q = 0Å^-1^ (I(0)), derived using the Guinier approximation I(q) = I(0) exp(-q2RG2/3) with the limits qRg <1.3). I(0) and RG values were compared for each collected SAXS curve (frame) across the entire elution peak (Fig. S4A). The elution peak was mapped by plotting the scattering intensity at q=0 Å^-1^ (I(0)), relative to the recorded frame. Uniform RG values across an elution peak represented a homogenous assembly (Fig. S4A). The merged experimental SAXS data were additionally investigated for aggregation by inspecting Guinier plots (Fig. S4*B*). The program SCÅTTER 4.0 was used to compute the distance distribution, P(r) (Fig. S4*E*). The distance r where P(r) approaches zero intensity is termed the D_max_ of the molecule. P(r) was normalized based on molecular mass (142kDa for the XPA-RPA-DNA FL complex and 131kDa for XPA_1-239_-RPAΔ70NΔ32N-DNA complex) determined by SAXS of assemblies as calculated by SCÅTTER as described _33_. The differences in the scattering power of protein and DNA were not taken into account in the determination of molecular mass because the contribution of DNA to the total is much less than that of protein. The SAXS data will be deposited to the SASBDB databank for the complexes with the 3’ junction and 5’ junction substrates.

### Computational Modeling

The structure of the XPA_1-239_/DNA/RPAΔ32NΔ70N complex was generated in four steps:

1. Two different initial models of the XPA_98-239_/RPA DNA Binding Core (RPA70AB + RPA70C/32D/14) complex bound to a model DNA substrate were prepared by using previously determined integrative structures^24,34^. The substrate consisted of a Y-shaped DNA with a 10-bp duplex, a 8-nt 5’ overhang and a 30-nt 3’overhang. Model A was generated by combining (i) a jointly refined SAXS and molecular dynamics simulation structure of RPA DBC bound to a 30-nt ssDNA substrate ^10^ and (ii) a model of XPA_98-239_/RPA70AB complex bound to a 3’ ss-ds DNA junction generated from a combination of homology modeling, NMR, SAXS, and molecular docking (22). RPA70AB is contained in both, so the structures could be aligned by superposing them on these two domains. Model B was generated using FoXS Dock by combining the same integrative model of the XPA_98-239_/RPA70AB/DNA complex with the RPA timer core (RPA70C/RPA32D/RPA14) without any pre-alignment.
2. A homology model of RPA32C bound to XPA_29-46_ was prepared using the NMR structure of RPA32C bound to XPA homologue UNG2 (PDB: 1dpu) as a template in Modeller 9v4 ^35^.
3. FoXS Dock was used to dock the structures from steps 1 and 2 using the SAXS profile of the full XPA_1-239_/DNA/RPAΔ32NΔ70N complex ^36,37^. A total of 4878 models were generated and the top model selected had the best combined SAXS χ and statistical potential scores. Although the results obtained were effectively the same for the two different initial models generated in step 1, the structural models used for further refinement were derived from initial model A.
4. Missing residues in XPA (N-terminal residues 1-28 and residues 47-97) and the linker between RPA32D and RPA32C were built using the program Modeller 9v4 ^38^. Then the FoXS server was used to best fit the model to the experimental SAXS profile.

The same strategy was used to generate the structural model of the complex with the Y-shaped DNA substrate composed of a 10-bp duplex, a 30-nt 5’ overhang and a 8-nt 3’overhang. In this case, in step 3 a better fit to the data was obtained for structural models derived from initial model B.

Visualization and superposition of models into the molecular envelopes generated by DENSS _39_ used the Chimera Fit to Map tool ^40^ and DENSS align ^39^.

### Lentiviral Cell Line Transduction

SV40-transformed human fibroblasts XP2OS (XPA mutant) were cultured in Dulbecco’s Modified Eagle’s Medium (DMEM, Cytiva) supplemented with 10% fetal bovine serum (FBS, Millipore) and penicillin streptomycin (P/S, Gibco) at 37°C in the presence of 5% CO2. For lentiviral transduction, 0.75 µg of pWPXL expression vector containing XPA cDNA, 2.25 µg of pMD2.G envelop plasmid, and 2.25 µg of psPAX2 packaging plasmid were transfected into 293T cells using Lipofectamine 3000 (L3000001, Thermo Fisher Scientific) following established protocols and virus harvested after 1 days ^41^. One day before transfection, XP2OS cells were seeded in 6-well plates at 50% confluency and incubated with lentivirus with a multiplicity of infection (MOI) of 2 for 24hr and subsequently grown as described above.

### Western blot

Cells were lysed with RIPA buffer (150 nM NaCl, 50 mM Tris-HCl (pH 8.0), 5 mM EDTA, 1% Triton X-100, 0.1% SDS, 0.05 g/mL sodium deoxycholate, 1 mM phenylmethylsulfonyl fluoride (PMSF), 5 mM MgCl_2_, supplemented with protease inhibitor (Roche)). Protein concentration was measured by Bradford assays (Thermo Fisher Scientific). After addition of LDS sample buffer containing 2.5% of 2-mercaptoethanol (4x, Invitrogen), the samples were heated to 95 °C for 10 min. Samples containing 25 µg of protein were separated in 8-16% Tris-Glycine gels (Thermo Fisher Scientific) at 150 V for 45 min, transferred to PVDF membrane (Amersham Hybond 0.2 um PVDF) at 100 V for 1hr 10 min with mini-protean tetra system (Bio-Rad) at 4 °C, and blocked with 5% skim milk in Tris-buffered saline containing 0.1% Tween-20 (TBST) for 1 hr at room temperature (RT). Rabbit polyclonal XPA antibodies (1:1,000), HA-tag antibodies (1:7,000), or Ku80 antibodies (1:3,000) were added to the TBST and incubated overnight at 4 °C. The blot was then incubated with anti-goat IgG rabbit antibody (1:3,000) for 1hr at RT and the bands visualized with chemiluminescent substrate (cat #: 34577, Thermo Fisher Scientific) and ChemiDoc touch imaging system (cat #: 1708370, Bio-Rad).

### Clonogenic cell survival assay

Native and transduced XP2OS cells were cultured in DMEM supplemented with 10% FBS and 1% P/S. 2000 cells were seeded in a 6 cm dish 1 day before UV irradiation. Cells were treated with 0, 1, 2, 4 J/m^2^ of UV-C, then cells were grown for 10 days. Cells were fixed with 4% paraformaldehyde for 15 minutes and stained with 1% of methylene blue for 2 hrs. After washing with water, colonies containing more than 25 cells were counted. The survival rate was normalized to the number of colonies of UV non-treated cells.

### Local UV irradiation assay

Cells were seeded onto slide cover glass (Neuvitro, cat# GG-22-PDL) 1 day before UV treatment and incubated with growth medium. For UV treatment, the medium was removed and cells were covered with 5 µm isopore membrane (cat #: TMTP04700, Merck) and irradiated with 100 J/m^2^ of UV-C. After UV irradiation, cells were incubated for different repair times (0, 1, 2, 4, 8, 24 hrs for (6-4) PPs repair assay/ 0, 8, 24, 48 hrs for CPDs repair assay). After washing with DPBS, cells were fixed with 4% paraformaldehyde in DPBS for 10 min at RT and permeabilized with DPBS containing 0.5% Triton X-100 for 5 min on ice. For CPDs and (6-4) PPs detection, DNA was denatured with 2 M HCl for 30 min at RT. Following a 30 min blocking step with DPBS containing 20% FBS, cells were incubated with diluted primary antibody in 5% FBS for 2 hrs at RT. Following 5 washing steps with DPBS, the cells were incubated with diluted secondary antibody in 5% FBS for 1 hr at RT and stained with DAPI (Vectashield with DAPI, cat #: H-1200-10).

### Co-localization assay

Cells were seeded and irradiated with UV-C as described for the local UV irradiation assay. After UV treatment, cells were lysed with cold hypotonic buffer (10 mM Tris-HCl (pH 8.0), 2.5 mM MgCl_2_, 10mM β-glycrophosphate, 0.2% Igepal, 0.2 mM PMSF, 0.1 mM Na_3_VO_4_) for 15 min on ice and washed with hypotonic buffer without Igepal for 4 min at RT. Cells were fixed in 2% formaldehyde in DPBS for 5 min at RT. After washing twice with 0.1% Triton X-100 in DPBS for 5min, cells were stored in 70% ethanol at -20 °C. After removal of the ethanol, cells were washed with DNAseI digestion buffer (10mM Tris-HCl (pH 8.0), 5 mM MgCl_2_) for 2 min and then the DNA digested with 20U DNaseI (cat #: D4527, Sigma-Aldrich) in 1 mL of digestion buffer for 40 sec at RT. EDTA (500 mM) was added to a final concentration is 10 mM EDTA to stop the reaction. The solution was removed and cells were washed twice with 10 mM EDTA in DPBS for 5min. After treatment with blocking buffer (1% BSA, 0.2% Tween-20), cells were incubated with diluted primary antibody (CPD (1:1,500) and HA-tag (1:250)) in blocking buffer for 2 hrs. Following 3 washing steps with DPBS containing 0.2% Tween-20, cells were incubated with diluted secondary antibody in blocking buffer and stained with DAPI.

### Slot-blot assay

Cells were irradiated with 5 J/m^2^ of UV-C and collected after different repair time (0, 2, 4, 8 hrs for (6-4) PPs and 0, 8, 24, 48 hrs for CPDs). Genomic DNA was prepared from harvested cells using QIAamp DNA mini kit (Qiagen) according to the manufacturer’s protocol. After adjusting genomic DNA concentration, DNA was denatured with 7.8 mM EDTA (for (6-4) PPs) or with 0.4 M NaOH and 10 mM EDTA (for CPDs). The denatured DNA was boiled at 95 °C for 10 min. DNA samples (200 ng of (6-4) PPs or 100 ng of CPDs) were neutralized by adding an equal volume of 2 M ammonium acetate (pH 7.0) and vacuum-transferred to a pre-washed nitrocellulose membrane using a BioDot SF microfiltration apparatus (Bio-Rad). Each well was washed 2 times with SSC buffer. The membrane was removed from the apparatus rinsed twice with SSC, air-dried, baked under vacuum at 80 °C for 2 hrs and blocked with 5% skim milk in PBS. For lesion detection, the membrane was incubated with (6-4) PPs (1:2,000) or CPDs antibodies (1:3,000) at 4 °C overnight and then incubated with anti-goat IgG mouse antibody (1:2,500 for (6-4) PPs or 1:5,000 for CPDs) for 1 hr at RT. The blot was visualized with ECL system (Thermo Fisher Scientific) and the total amount of DNA loaded on the membrane was visualized with SYBR-gold (Thermo Fisher Scientific) staining.

### In vitro NER activity assay with cell extracts

A plasmid containing a site-specific acetylaminofluorene (AAF) lesion was incubated with XPA-deficient (XP2OS) cell extract in the absence or presence of purified wild-type or mutant XPA proteins ^25^. For each reaction, 2 μl of repair buffer (200 mM HEPES-KOH, 25 mM MgCl_2_, 110 mM phosphocreatine (di-Tris salt, Sigma), 10 mM ATP, 2.5 mM DTT and 1.8 mg/ml BSA, adjusted to pH 7.8), 0.2 μl of creatine phosphokinase (2.5 mg/ml, Sigma), 3 μl of XPA-deficient cell extract (about 10 mg/ml), NaCl (to a final concentration of 70 mM) and 50 nM of purified wild-type or mutant XPA in a total volume of 9 μl were pre-warmed at 30 °C for 10 min. 1 μl of plasmid containing AAF (25 ng/μl) was added to reaction mixture and incubated at 30 °C for different incubation times (0, 5, 10, 20, 45, and 90 min). 0.5 μl of 1 μM of a 3′-phophorylated oligonucleotide for product labeling was added and the mixture heated at 95 °C for 5 min. The mixture was allowed to cool down to room temperature for 15 min. 1.2 μl of a Sequenase/[α-^32^P-dCTP mix (0.25 units of Sequenase and 2.5 μCi of [α-^32^P]-dCTP per reaction) was added and the mixture incubated at 37 °C for 3 min. Then 1.2 μl of dNTP mix (100 μM of each dATP, dTTP, dGTP; 50 μM dCTP) added to mixture and incubated for another 12 min. The reactions were stopped by adding 12 μl of loading dye (80% formamide/10 mM EDTA) and heating at 95 °C for 5 min. 6 ul of sample was loaded on 14% sequencing gel (7 M urea, 0.5x TBE) at 45 W for 2 hrs 30 min. The reaction products were visualized using a PhosphorImager (Amersham Typhoon RGB, GE Healthcare Bio-Sciences). Two independent repetitions were performed. The NER products were quantified using ImageQuant TL and normalized to the amount of NER product formed with WT-XPA at 90 min.

### In vitro NER activity assay with purified proteins

The same protocol was used as for the assays with cell extracts, except that purified NER proteins were used. For each reaction, 5 nM of XPC-RAD23B, 10 nM of TFIIH, 20 nM of XPA, 41.6 nM of RPA, 27 nM of XPG, and 13.3 nM of XPF-ERCC1 was used. All proteins were >95% pure and produced as previously described: XPC-RAD23B ^42^; TFIIH ^43^; RPA ^44^; XPG ^45^; ERCC1-XPF ^46^.

## Results

### XPA mutations inhibit binding to RPA

XPA has been shown to interact with RPA32C via disordered residues 29-46 and with RPA70AB through the zinc binding motif in the DBD ^20,24^. We set out to inhibit the interaction between XPA and RPA and assess the importance of these interactions for NER using site-specific mutagenesis to generate a series of XPA mutations in these two regions (**Fig. 1A, Table 1**). We first assessed the two interaction interfaces individually and then in combination.

**Table 1.** Mutant XPA proteins used in this study.

| **Name** | **Mutation** | **Binding site** |
| --- | --- | --- |
| **XPA 32-E1** | R34E | 32C |
| **XPA 32-E3** | R34E/R39E/R42E | 32C |
| **XPA 32-E4** | R30E/K31E/R32E/R34E | 32C |
| **XPA 32-E6** | R30E/K31E/R32E/R34E/R39E/R42E | 32C |
| **XPA 70-M1K** | E106K | 70AB |
| **XPA 70-M1A** | F112A | 70AB |
| **XPA 70-M2** | E106K/F112A | 70AB |
| **XPA 70-M4** | D101N/E106K/K110E/F112A | 70AB |
| **XPA 70-M6** | D101N/E106K/K110E/E111K/F112A/D114N | 70AB |
| **XPA 32-E1/70-M1A** | R34E/F112A | 32C &70AB |
| **XPA 32-E1/70-M2** | R34E/E106K/F112A | 32C &70AB |
| **XPA 32-E3/70-M2** | R34E/R39E/R42E/ E106K/F112A | 32C &70AB |
| **XPA 32-E4/70-M2** | R30E/K31E/R32E/ R34E/E106K/F112A | 32C &70AB |

**Figure 1:**
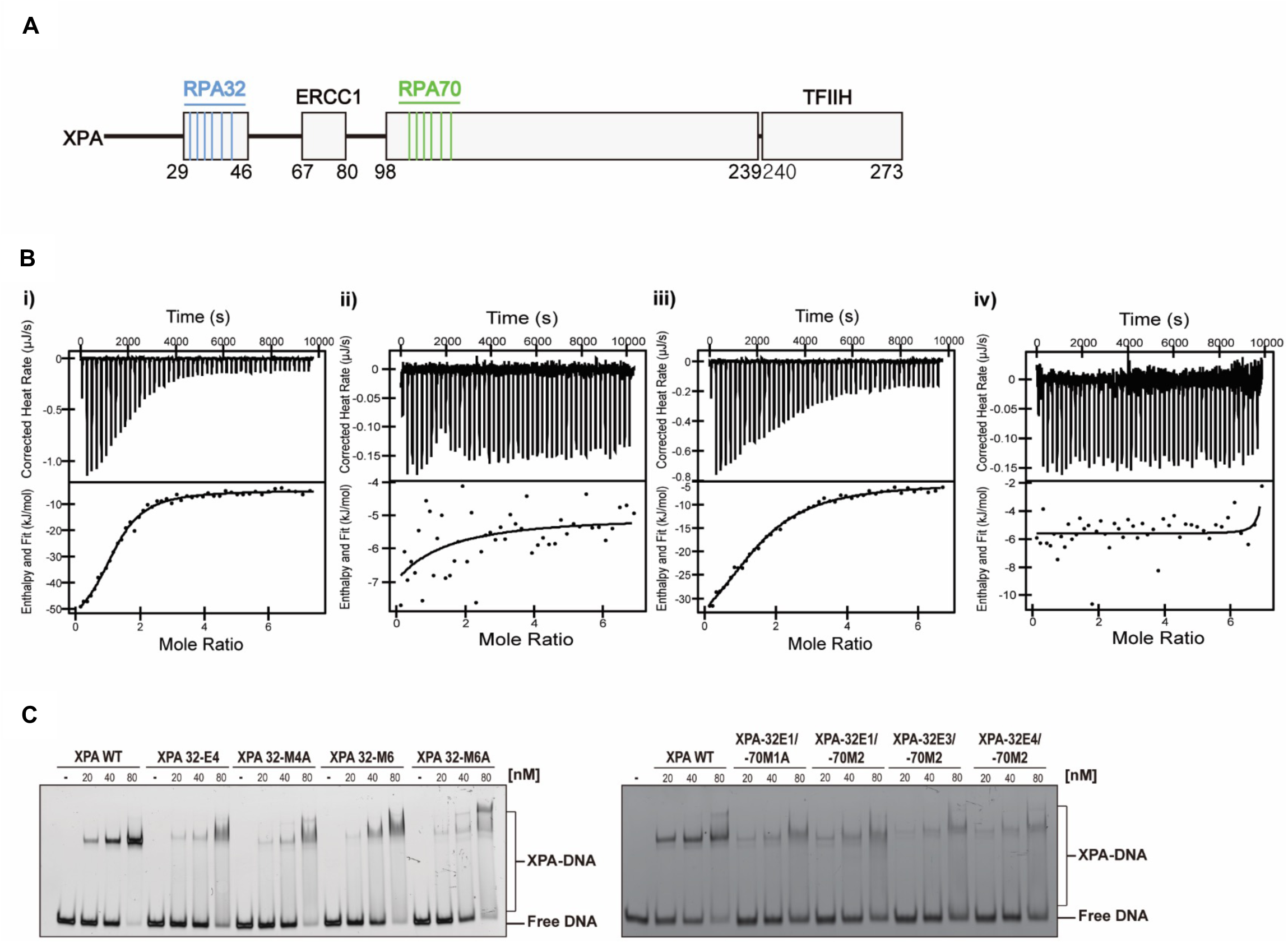
Interaction of XPA wild-type, XPA-32-4E, XPA-70-M2 and XPA-32-4E/70-M2 with RPA. **A**. The domain map of the XPA protein with interaction sites with other protein and mutants introduced in this study highlighted. **B**. Isothermal titration calorimetry thermograms of XPA mutants and full-length RPA showing the raw heat release (*top*) and integrated heat release (*bottom*). Titrations of RPA were performed with i) wild-type XPA, ii) XPA 32-E4, iii) XPA 70-M2, and iv) XPA 32-E4/70-M2. For the titrations in i)-iv), the first injection of 1 µL was removed for analysis. The thermograms in i-iv are representative of one of three or more replicates. **C**. EMSA analysis of XPA proteins with mutation in the RPA32 and RPA30/70 interaction domains binding to a 5’-FAM-labeled three-way DNA junction (100nM). Protein concentrations and position of DNA on the gel are indicated

We have previously mapped the interaction of XPA_29-46_ to an acidic binding surface on RPA32C ^20^. There is a strong electrostatic component to this interaction, mediated by six basic XPA residues: Arg30, Lys31, Arg32, Arg34, Arg39, and Arg42. To test their role in mediating the interaction with RPA32C, a series of charge reversal mutations of these residues to Glu were made to obtain the strongest possible inhibitory effect **(Table 1)**. We have previously identified residues Asp101, Glu106, Lys110, Glu111, Phe112, and Asp114 of XPA as important in mediating the interaction with RPA70AB and have shown that mutations in these residues diminish the interaction between XPA and RPA70 and inhibit the NER reaction *in vitro* ^24^. These same mutations were used here for the cell-based assays, both alone and in combination with the RPA32C mutations. All XPA mutants used in this study are shown in **Table 1**.

To validate the mutant design, we tested the ability of the mutant XPA proteins to interact with RPA and DNA. After purifying the recombinant proteins (**Fig. S1A**), the affinity of RPA for XPA wild type and variants with mutations in the RPA32C and RPA70AB binding motifs was measured using isothermal titration calorimetry (ITC). A dissociation constant (K_d_) of 3.6 ± 1.2 µM was determined for the wild type protein. Whereas mutations in the RPA32C binding motif had a dramatic effect on the binding of XPA to RPA, mutations in the RPA70AB binding motif had a much more limited effect, no great than a 3-fold reduction in affinity compared to the wild-type protein (**Fig. 1B, Table 2**). The loss of binding between RPA and the 32C-4E mutant, independent of a mutation in the RPA70AB binding motif, supports the previous finding that the RPA32C-XPA_29-46_ interaction is stronger than the interaction between RPA70AB and XPA_98-239_ ^20,24^.

**Table 2.**
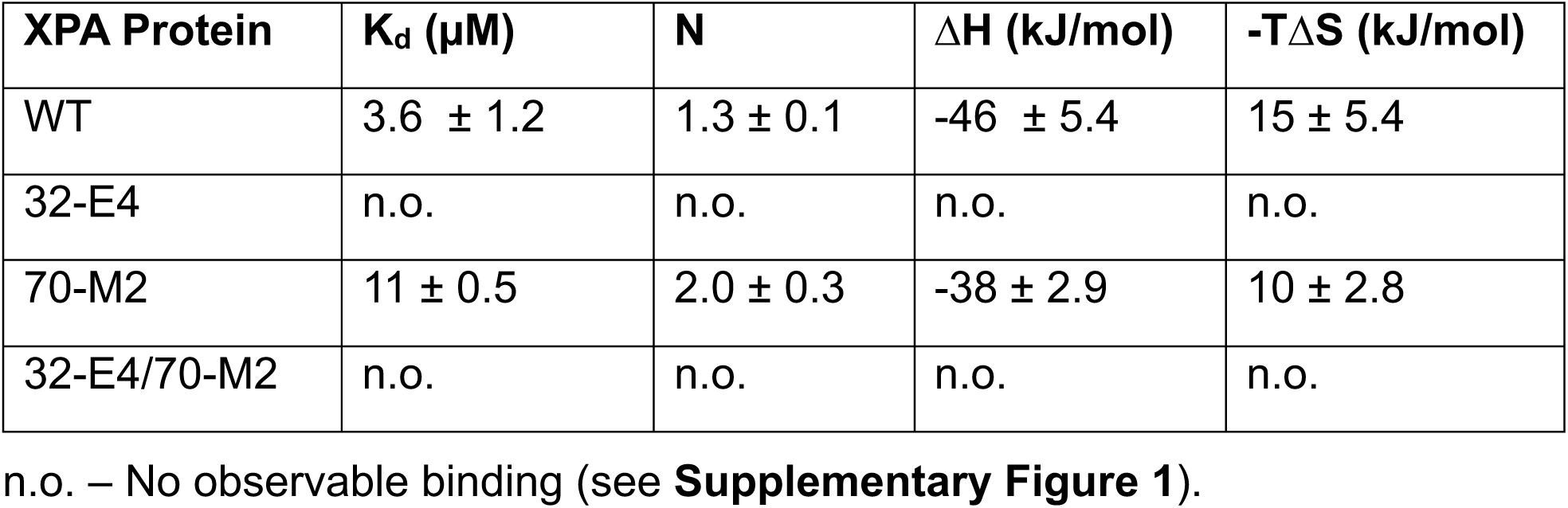
Thermodynamic parameters for the binding of XPA to RPA.

| **XPA Protein** | **K<sub>d</sub> (μM)** | **N** | **ΔH (kJ/mol)** | **-TΔS (kJ/mol)** |
| --- | --- | --- | --- | --- |
| WT | 3.6 ± 1.2 | 1.3 ± 0.1 | -46 ± 5.4 | 15 ± 5.4 |
| 32-E4 | n.o. | n.o. | n.o. | n.o. |
| 70-M2 | 11 ± 0.5 | 2.0 ± 0.3 | -38 ± 2.9 | 10 ± 2.8 |
| 32-E4/70-M2 | n.o. | n.o. | n.o. | n.o. |

n.o. – No observable binding (see **Supplementary Figure 1**).

We next tested the ability of the XPA mutants to bind DNA substrates mimicking NER intermediates using electric mobility shift assays (EMSA) (**Fig. 1C**). We have previously shown that mutations in the RPA70AB binding interface do not significantly affect protein folding or DNA binding ability ^24^. A similar observation was made for the XPA proteins with mutations in the RPA32C binding motif, as EMSA measurements showed their DNA binding affinity was comparable to WT-XPA (**Fig. 1C**). We note that the shape of the DNA-bound bands was slightly different, possibly due to the presence of additional negatively charged residues in the protein. Proteins with mutations in both the RPA32C and RPA70AB interaction interfaces also showed robust DNA binding activity, albeit with about 2-fold lower binding affinity than WT protein (**Fig. 1C**).

To further evaluate any perturbations of XPA binding to the DNA substrate, we quantified DNA binding affinity by microscale thermophoresis (MST) using the fluorescently tagged NER junction mimic reported previously ^47^. This approach was used to measure the affinity of the 70-M2 mutant in the XPA DNA binding domain (DBD) construct ^24^ and saw had no discernable effect. A similar observation was made here for the XPA full-length mutant protein. In addition, as anticipated based on their location in the XPA N-terminal region distant from the DBD, very little effect on DNA binding affinity was observed for the 32-4E mutation (K_d_ = 8.4 ± 2.3 µM versus 5.4 ± 1.1 µM for the WT, respectively; **Fig. S1B**). In the construct combining the 32-4E and 70-M2 mutations, there was a slightly larger effect (K_d_ = 10 ± 3 µM). We are unsure of the origin of these very modest effects on binding of DNA, but they are clearly much smaller than the many orders of magnitude of the effect on XPA-RPA interaction caused by mutations in the RPA32C binding motif.

### Inhibition of XPA interaction with RPA70AB reduces UV-induced damage repair

We have previously shown that the mutations in the RPA70AB binding motif of XPA inhibit the physical interaction and reduce the biochemical NER activity on a lesion-containing plasmid ^24^. To investigate the effect in cells, we generated stable XP2OS cell lines expressing XPA with mutations in the RPA70AB interaction domain by lentiviral transfections. We verified that these cells expressed WT and mutant XPA proteins at comparable levels (**Fig. 2A**). The cells were first tested for hypersensitivity to UV irradiation. Compared to WT-XPA, XPA 70-M1K, 70-M1A, and 70-M2 showed a slight, but statistically significant increase in UV sensitivity (**Fig. 2B**). Cells expressing XPA 70-M4 and M6 with a larger number of mutations exhibited a further increase in UV sensitivity (**Fig. 2B**).

**Figure 2:**
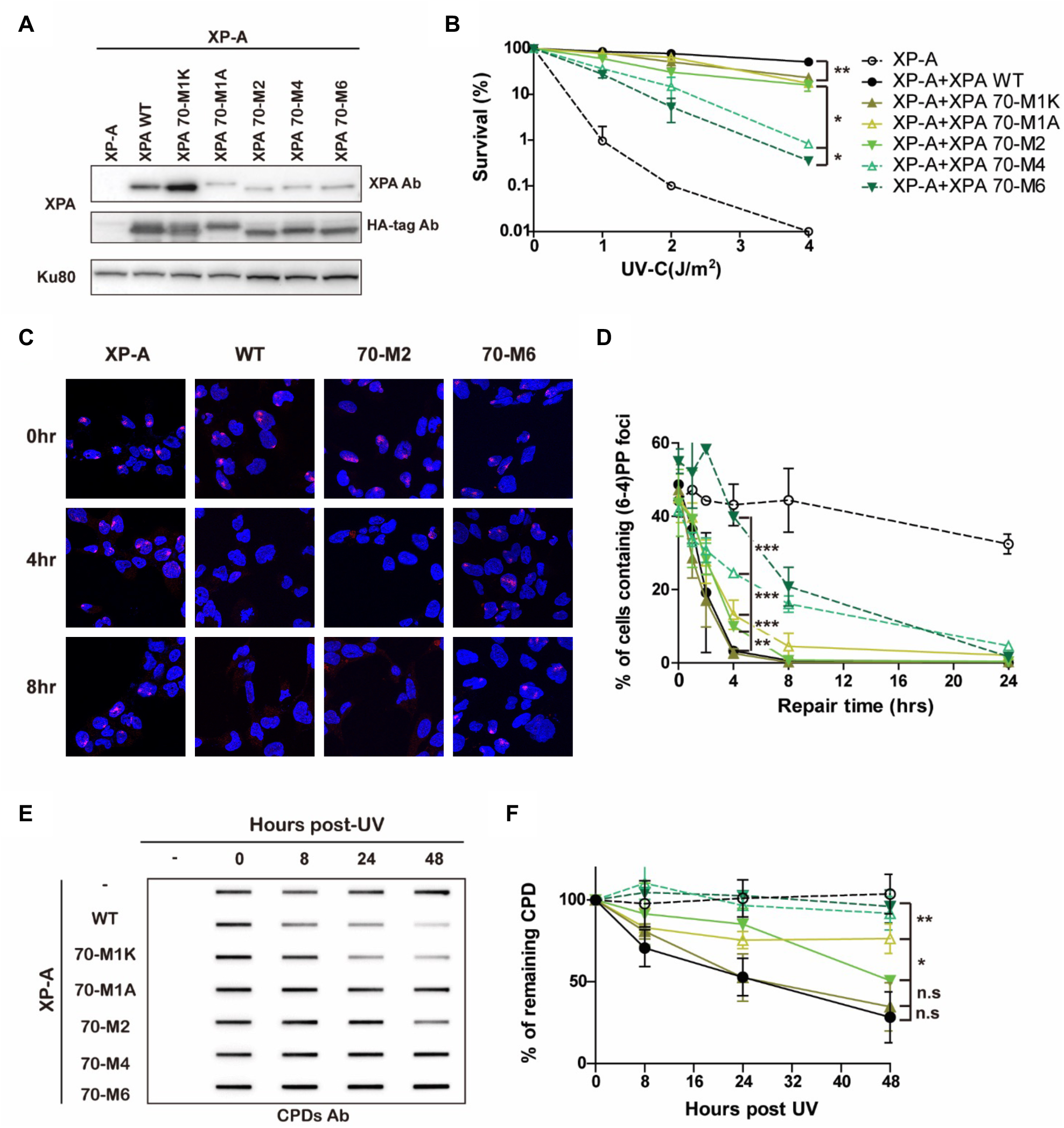
Mutations in the RPA70 interaction domain of XPA cause defects in the repair of UV lesions. **A**. Expression level of WT and RPA70-interaction mutant XPA in XP2OS cells transduced with HA-tagged XPA. Protein detected with anti-XPA and anti-HA antibodies, using Ku80 as a loading control. **B**. Clonogenic survival assays. Cells were treated with 0, 1, 2, 4 J/m^2^ of UV irradiation, grown for 10 days and stained with methylene blue. Survival rates were normalized to the colony number of non-treated cells. The p-value was measured compared to XPA WT. \**P*< 0.05, \*\**P*<0.01, \*\*\**P*<0.001. **C**. Representative figures of cells irradiated through a 5 µm micropore filter with UV (100 J/m^2^) and stained with a (6-4) PP antibody after the indicated times of repair. (6-4) PP foci are red and cell nuclei are stained blue with DAPI. **D**. Quantification of (C). 100 cells were counted for each sample and the data represent at least 2 independent experiments. The p-value was measured compared to XPA WT. \**P*< 0.05, \*\**P*<0.01, \*\*\**P*<0.001. **E**. Determination of CPD repair kinetics using slot-blot assays. Cells were irradiated with 5 J/m^2^, genomic DNA isolated at indicated time points and the adduct levels determined using an anti-CPD antibody CAC-NM-DND-002. **F**. Quantification of E. Band intensities were normalized to the WT band at 0 hr. The p-value was measured compared to XPA WT. \**P*< 0.05, \*\**P*<0.01, \*\*\**P*<0.001.

To obtain deeper insights, we examined the repair kinetics of the UV lesions 6-4 photoproducts ((6-4) PPs) and cyclopyrimidine dimers (CPDs) in these cells by: (i) measuring the adduct levels at sites of local UV irradiation in cell nuclei assay and (ii) measuring the level of adducts in globally irradiated cells using slot-blot assays. Following irradiation through micropore filters, 40∼60% of cells contained sites of (6-4) PP damage. In XP-A deficient cells, the (6-4) PPs persisted for 24 hrs post UV irradiation, whereas these adducts were repaired within 4 hrs in WT-XPA cells (**Fig. 2C, D**). The kinetics of (6-4) PP repair corresponded well to the results of survival assays: XPA 70-M1K exhibited WT kinetics of repair, XPA 70-M1A and XPA-70M2 had slightly slower repair kinetics, and XPA-70M4 and XPA-70M6 had the slowest repair kinetics although the (6-4) PPs were still repaired at much higher levels than in XPA deficient cells (**Fig. 2B-D**). The determination of repair rates using slot blot assays with (6-4) PP antibodies showed similar results while allowing for better quantification (**Fig. S2A, B**). The repair of (6-4) PPs was near WT levels in XPA 70-M1K, reduced by about 2-fold in XPA 70-M1A and XPA 70-M2, and by about 5-fold in XPA-70M4 and XPA-70M6 cells. The effects of the mutations were also measured for CPD repair kinetics, which are known to be repaired significantly more slowly than (6-4) PPs. The differences in CPD repair rates paralleled those observed for (6-4) PPs. The results from the repair of the CPDs at sites of local UV damage were more difficult to quantify, due to the slower repair rates leading to foci with gradually decreasing intensity (**Fig. S2E** and **S2F**). However, data from the slot bots of the globally irradiated cells revealed an ∼2-fold decrease in CPD repair for XPA-F112A and XPA-M2 and an ∼5-fold decrease for XPA-70M4 and XPA-70M6 (**Fig. 2E, F, S2C-F**). The CPD repair rates for XPA-70M4 and XPA-70M6 were statistically different from wild-type cells (**Fig. 2E, F, S2C, D)**. Together our data indicate that progressive weakening of the XPA-RPA70 interaction leads to hypersensitivity to UV irradiation due to reduced rates of NER.

### Interaction between XPA and RPA32C is required for progression of NER

We took an equivalent cell-based approach to assesses the importance of the interaction of XPA with RPA32C for the repair of UV lesions by NER. We generated cells with mutations in the RPA32C-interaction motif of XPA (**Table 1**) and picked clones with comparable XPA expression levels (**Fig. 3A**). Then their sensitivity to UV irradiation and lesion repair kinetics was tested. Following UV treatment, XPA32-E1 cells showed similar sensitivity as WT-XPA cells (**Fig. 3B**), while the XPA32-E3, 32-E4, and 32-E6 cells showed greater, statistically significant hypersensitivity to UV, although they were less sensitive than XP-A deficient cells (**Fig. 3B**). Survival rate of XPA 32-E3, 32-E4, and 32-E6 were not statistically different from each other. We next investigated the repair of UV-induced (6-4) PPs and CPDs of the mutant cells by local UV irradiation assays and slot-blot assays. The results were consistent between the two assays, with the XPA 32-E1 single mutant showing similar levels of (6-4) PP repair to XPA-WT, while XPA 32-E3, 32-E4, and 32-E6 showed a 3-6 fold decrease in the rate of repair (**Fig. 3C, D, S3A, B**). Similarly, there was not difference in the repair of CPDs between XPA-WT and XPA 32-E1, whereas CPDs persisted in XPA 32-E3, 32-E4, and 32-E6 up to 48 hrs at levels similar to XPA-deficient XP2OS cells (**Fig. 3E, F, S3C-F**). In conclusion, a progression in the degree of defect in the interaction between XPA and RPA32C or RPA70AB led to increasingly reduced but still significant NER activity and ability to repair UV-induced damage.

**Figure 3:**
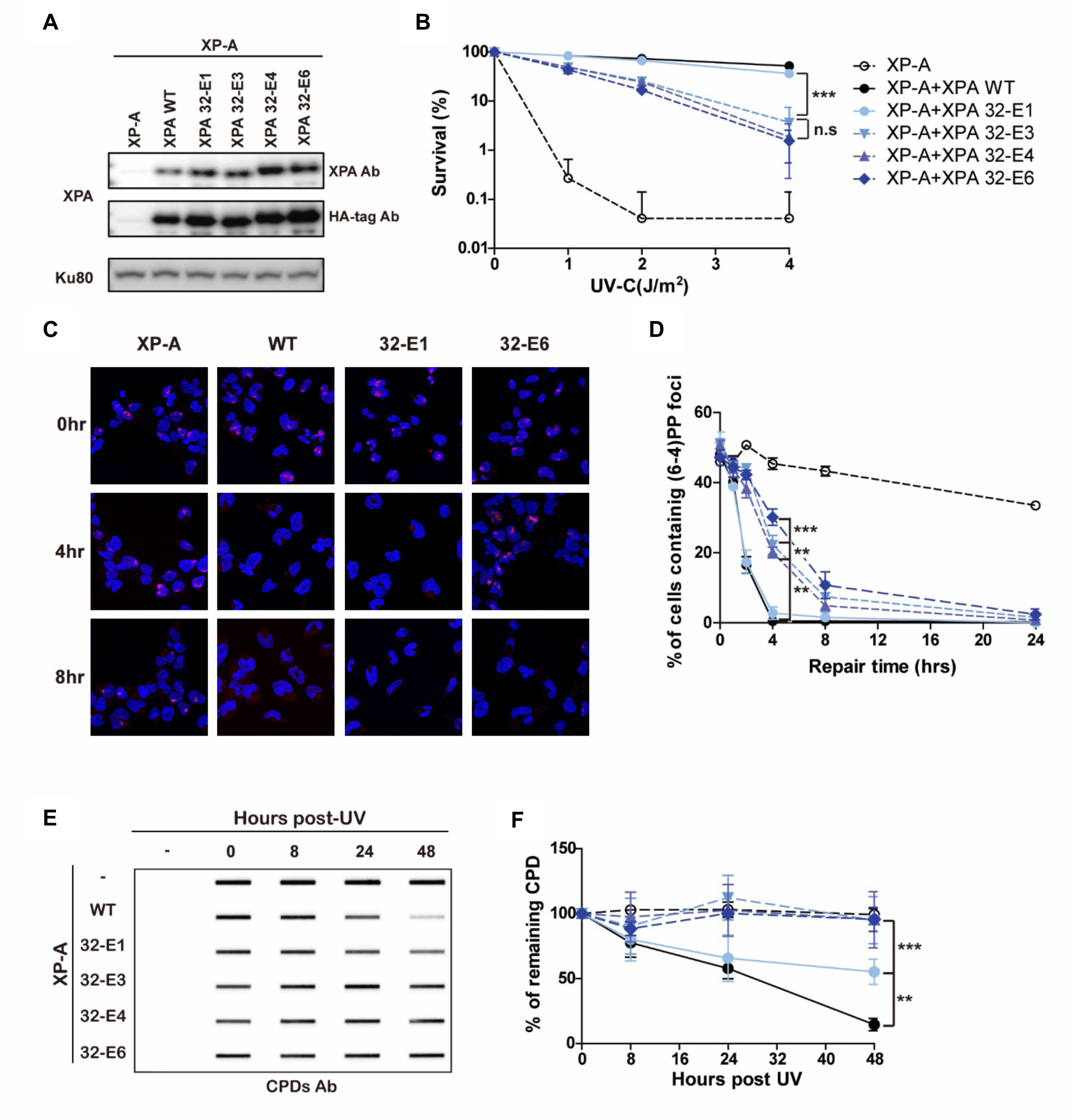
Mutations in the RPA32 interacting domain of XPA lead to a cellular defect in the repair of UV lesions. **A**. Expression level of WT and RPA32-interaction mutant XPA in XP2OS cells transduced with HA-tagged XPA. Proteins detected anti-XPA and anti-HA antibodies, using Ku80 as a loading control. **B**. Clonogenic survival assays. Cells were treated with 0, 1, 2, 4 J/m^2^ of UV irradiation, grown for 10 days and stained with methylene blue. Survival rates were normalized to the colony number of non-treated cells. \**P*< 0.05, \*\**P*<0.01, \*\*\**P*<0.001 **C**. Representative figures of cells irradiated through a 5 µm micropore filter with UV (100 J/m^2^) and stained with a (6-4) PP antibody after indicated repair times. (6-4) PP foci are red and cell nuclei are stained blue with DAPI. **D**. Quantification of C. 100 cells were counted for each sample and the data represent at least 2 independent experiments. The p-value was measured compared to XPA WT. \**P*< 0.05, \*\**P*<0.01, \*\*\**P*<0.001. **E**. Determination of CPD repair kinetics using slot-blot assays. Cells were irradiated with 5 J/m^2^ genomic DNA isolated at the indicated time points and the adduct levels of genomic DNA determined with an anti-CPD antibody. **F**. Quantification of E. Band intensities were normalized to the WT band at 0 hr. The p-value was measured compared to XPA WT. \**P*< 0.05, \*\**P*<0.01, \*\*\**P*<0.001.

### Interactions of XPA with RPA32C and RPA70AB contribute synergistically to NER

Following the analysis of the individual contacts with RPA, we combined mutations in the RPA32C and RPA70AB binding interfaces of XPA to determine if loss of these physical interactions have independent or synergistic effects on NER activity and sensitivity to UV irradiation. To this end, we generated cells with mutations in both the RPA32C and RPA70AB interaction interfaces and compared their activities with the corresponding mutants in just one or the other interface (**Table 1, Fig. 4A**). We first tested the sensitivity of those cells to different doses of UV-C (**Fig. 4B**). Combining a limited number of mutations, such as XPA32-E1 and 70-M2 led to 2-fold reduction in sensitivity compared to individual XPA 70-M2, a clearly additive effect (**Fig. 4B**). The effect was even more dramatic when combining 70-M2 and 32-E3 or 32-E4 mutations. The difference in UV sensitivity of the XPA-32-E3/70-M2 and XPA-32-E4/70-M2 was no longer statistically significantly different from the XP2OS patients at a dose UV of 4 J/m^2^, while the corresponding individual 32-E3/E4 or 70-M2 mutations individually only showed moderate UV sensitivity (**Fig. 4B**).

**Figure 4:**
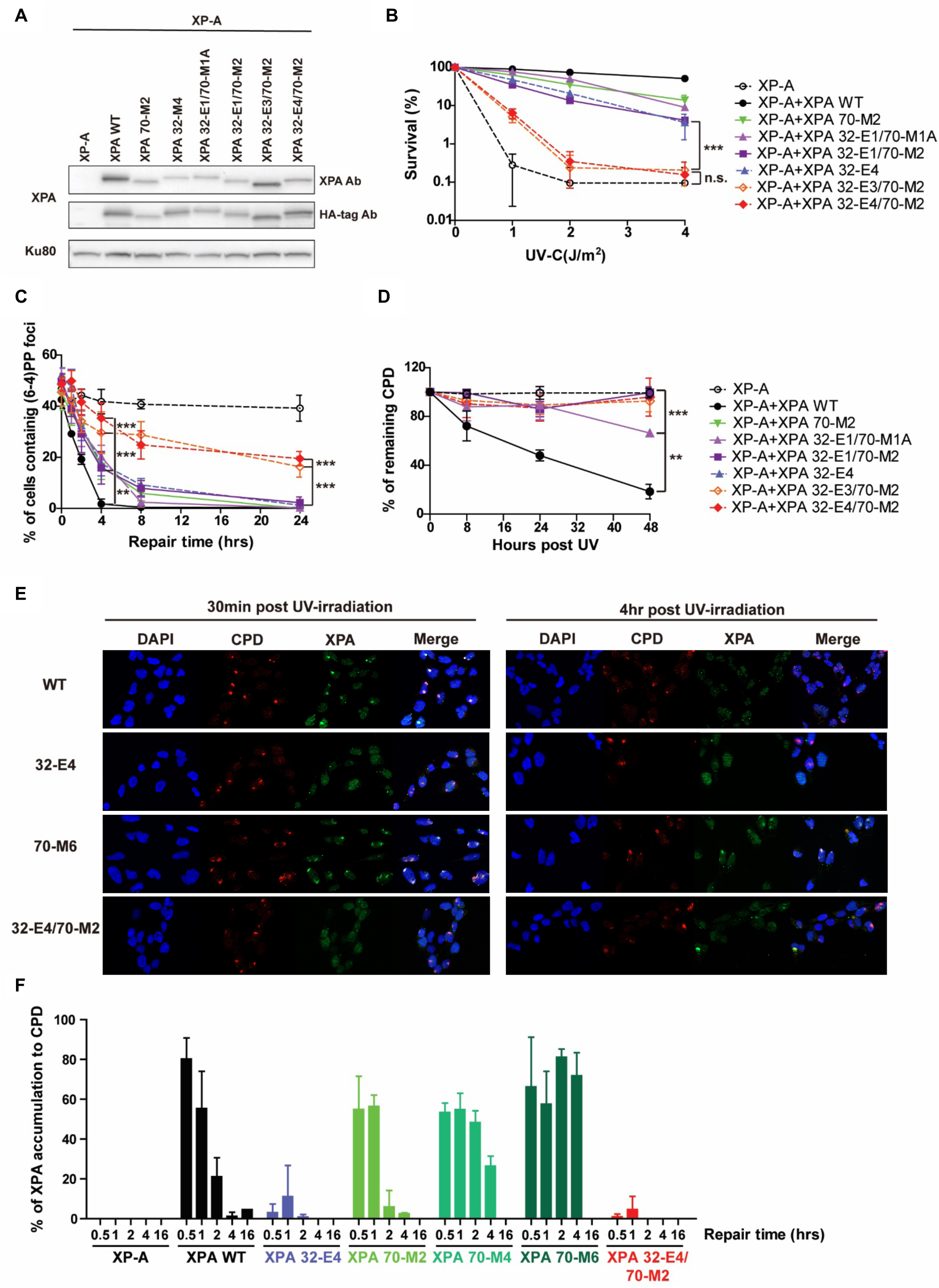
The RPA32 and 70 binding domains of XPA synergistically contribute to NER activity in cells. **A**. XPA expression levels in XP2OS cells transduced with HA-tagged WT and RPA32/70-interaction mutant XPA detected anti-XPA and anti-HA antibodies, using Ku80 as a loading control. **B**. Clonogenic survival assays. Cells were treated with 0, 1, 2, 4J/m^2^ of UV irradiation, grown for 10 days and stained with methylene blue. Survival rates were normalized to the colony number of non-treated cells. The p-value was measured compared to XPA 32-M4. \**P*< 0.05, \*\**P*<0.01, \*\*\**P*<0.001. **C**. Quantification of (6-4) PPs repair at sites of local UV damage. Cells were irradiated through a 5 µm micropore filter with UV (100 J/m^2^) and stained for (6-4) PP to measure repair kinetics. 100 cells were counted for each sample and the data represent at least 2 independent experiments. The p-value was measured compared to XPA WT. \**P*< 0.05, \*\**P*<0.01, \*\*\**P*<0.001. **D**. Determination of CPD repair kinetics using slot-blot assays. Cells were irradiated with 5 J/m^2^, genomic DNA isolated at the indicated time points and adduct levels of genomic DNA determined with an anti-CPD antibody. Band intensities were normalized to the intensity of WT band at 0 hr. The p-value was measured compared to XPA WT. \**P*< 0.05, \*\**P*<0.01, \*\*\**P*<0.001. **E**. Representative figure of co-localization assay with CPD and XPA. Cells were irradiated through 5 µm micropore filter with UV (100 J/m^2^) and stained for CPD and XPA to measure the accumulation of XPA to CPD. XPA was stained by HA-tag antibody. **F**. Quantification of E. 100 cells were counted for each sample and data represent 2 independent experiments. % of co-localization was calculated by dividing number of cells containing co-localization by number of cells containing CPD foci.

The progressive effect of cumulative mutations was also reflected in the repair kinetics of UV lesions. XPA 32-E1/70-M1A and XPA 32-E1/70-M2 cells exhibited slower repair of (6-4) PPs at the 8 hrs timepoint in both locally damaged sites and globally irradiated cells. A more dramatic loss in the ability to repair UV damage was observed for XPA 32-E3/70-M2 and XPA 32-E4/70-M2 (**Fig. 4C, S4A, B, E**). In these cells, up to 50% of (6-4) PPs remained at 24 hrs post UV, while 6-4PPs were never observed at these time points in any of the individual RPA-binding domain mutants. Indeed, in the slot blot assays the repair of (6-4) PPs in these cells were not statistically different from the XP2OS cells. The defects in repair kinetics were equally significant for CPDs. Whereas WT-XPA removed 80% of the CPDs within 48 hrs, these lesions persisted in cells expressing XPA with mutations in both RPA interaction domains to the same degree as in the XP-A patient cells (**Fig. 4D, S4C, D, F, G**). Together, these data clearly show that combining mutations in two XPA-RPA interaction interfaces are synergistic, leading to greater NER deficiency than either of the individual contacts alone.

Having shown that the RPA32C and RPA70AB interaction domains of XPA are important for NER, we aimed to determine what their respective roles might be. We therefore tested the kinetics of the arrival and departure of XPA with the XPA-RPA32C and XPA-RPA70AB at UV-induced DNA damage. In WT cells, the XPA protein arrived at UV damaged sites within 30 minutes and was released within 4 hrs (**Fig. 4E and 4F**). In the RPA32C interaction mutant XPA 32-E4 mutant cells, XPA localization to DNA damage was dramatically reduced (3% of UV-induced DNA damage at 0.5 hr and 10% of UV-induced DNA damage at 1 hr), suggesting that recruitment and/or association with the NER complex was diminished. By contrast, in cells expressing the RPA70AB interaction mutant, XPA associated with UV damage like the WT cells at 0.5 hr and 1 hr, but remained bound at damaged sites for extended periods of time (80% for 70-M6 and 50% for 70-M4 versus 20% for WT at 2hrs, 70% for 70-M6 and 25% for 70-M4 versus 1% of WT at 4hrs) (**Fig. 4F**). The XPA 32-E4/70-M2 double interaction domain mutant barely localized to UV damage, as expected from the XPA 32-E4 mutant **(Fig. 4E and F**). These results indicate that RPA32C and RPA70AB have distinct roles for XPA in NER: the RPA32C interaction to XPA is required for recruitment of XPA to UV-induced damage, while RPA70AB interaction to XPA is important for positioning of XPA for completing of NER, likely to license full complex assembly and 5’ incision by ERCC1-XPF.

### Mutations in XPA-RPA interaction domains inhibit NER in vitro activity

We next tested the intrinsic ability of the mutant XPA proteins to mediate the NER reaction using biochemical experiments by monitoring the excision of a damage-containing oligonucleotide from a plasmid containing a site-specific dG-AAF lesion using NER proficient cell extracts or the purified core NER proteins (XPC-RAD23B, TFIIH, XPA, RPA, ERCC1-XPF, and XPG) in the presence of WT and mutant XPA proteins. We first carried out the reaction using extracts from XPA-deficient XP2OS cell extracts supplemented with purified NER proteins. While incubation with extract alone did not yield a signal, we observed robust, time-dependent accumulation of the excision product in the presence of XPA WT and XPA32-E1 proteins (**Fig. 5A, lanes 1-13)**. By contrast, complementation with the XPA32-E4 protein led to an about 5-fold reduction in excision activity. (**Fig. 5A, lanes 14-19**). The activity of XPA 70-M1A and 70-M2 was decreased by about 2-fold (**Fig. 5A, lanes 20-31**). Combining these last two mutations with XPA32-E1, lead to a further reduction in activity (**Fig. 5A, lanes 32-43**). No excision activity was detectable in the presence of the XPA32-E4/70-M2 protein, consistent with the dramatic reduction observed in cellular activity (**Fig. 5A**, lanes 44-49).

**Figure 5:**
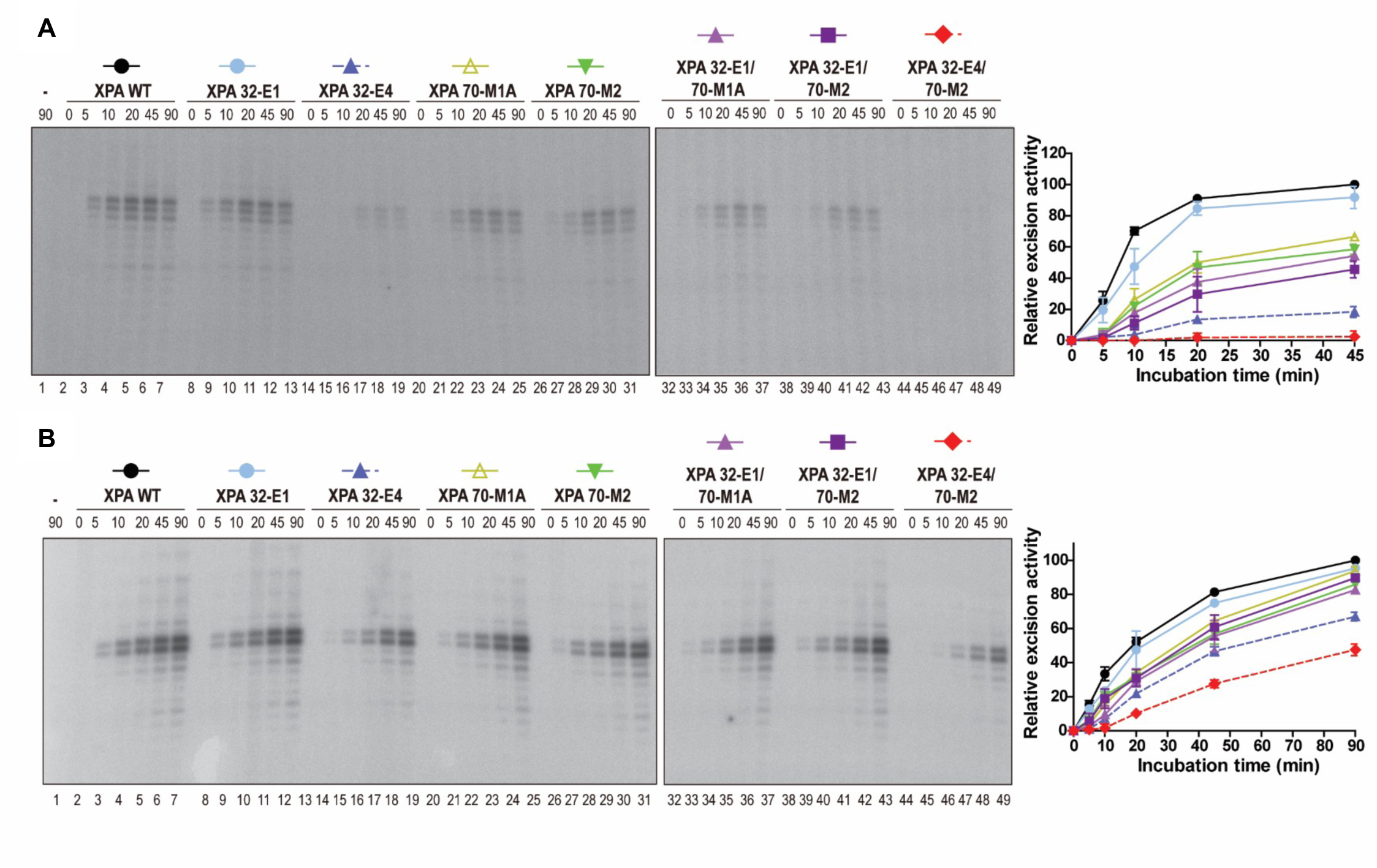
*In vitro* NER activity is diminished by mutations in the RPA32 and RPA70 interaction domains of XPA. **A**. Complementation of *in vitro* NER activity of XPA-deficient cell extracts with WT and RPA-interaction mutant XPA. A plasmid containing a site-specific AAF lesion was incubated with XP2OS cell extract and the purified XPA (50 nM) proteins for 0-90 min. The excision products were detected by annealing to a complementary oligonucleotide with a 4dG overhang, which was used as a template for a fill-in reaction with [α-^32^P] dCTP. Quantification of the data is from two independent experiments. **B**. *In vitro* NER activity of WT and RPA-interaction mutant XPA using purified NER proteins. Assays were conducted as in A., except that 20 nM XPA was used with purified XPC-RAD23B (5 nM), TFIIH (10 nM), RPA (42 nM), XPG (27 nM) and XPF-ERCC1 (13 nM) proteins were used instead of XP2OS cell extracts.

The overall NER activity was higher when purified proteins were used for the NER reaction, but the same order of activity level was observed for the various mutant versions of XPA (**Fig 5B**). The activity of the XPA 32-E4 protein was reduced by about 2-fold and for the XPA 32-E4/70-M2 protein by about 4-fold at the 45 minutes time point (**Fig. 5B, lanes 6, 18, 48**). In conclusion, the results from the biochemical experiments are in good agreement with the cellular studies and corroborate the observation the multiple weak interactions between XPA and both RPA32 and RPA70 are required for progression through the NER pathway.

### The two XPA-RPA contacts can be simultaneously accommodated

Having established that the interactions between XPA and RPA both affect NER proficiency, we considered whether they act either together or independently in promoting repair. One particularly relevant question in this context is whether both contacts can be engaged simultaneously or if they would need to be spaced in time along the NER trajectory? We therefore set out to determine the structure of the complex of XPA and RPA on model NER bubble substrates. As noted above, XPA and RPA are modular proteins with multiple globular and disordered domains and therefore are expected to retain considerable flexibility even when bound to each other. Such systems are not readily amenable to traditional high resolution structural techniques such as X-ray crystallography. However, in this case a lower resolution solution technique, small angle X-ray scattering (SAXS), is sufficient to provide the structural information of interest.

We have previously established that the XPA-DBD can form stable ternary complexes with RPA70AB and model substrates mimicking the ss-ds DNA junction of the NER bubble either 3’ or 5’ to the lesion ^24^. At first glance, binding to the 5’ junction poses a topological problem because engaging the ssDNA with 5’-3’ polarity (70A-70B-70C-32D) would seem to place RPA70AB far from the XPA DBD at the 5’ ss-ds junction. However, we found that RPA70AB formed similar complexes with XPA DBD by inverting its orientation to retain 5′-3′ polarity on the ssDNA. Thus, we continued to test if RPA-bound XPA is able to engage both 3’ and 5’ ss-dsDNA junction substrates.

Following the strategy used for study of the complex of XPA-DBD and RPA70AB bound to model substrates ^24^, we designed and optimized Y-shaped DNA substrates for both ss-ds junctions 3’ and 5’ to the lesion. These substrates were composed of dsDNA and a short ssDNA overhang for XPA and a longer ssDNA overhang for RPA. The optimal substrates contained a 10-bp duplex with one 8-nt and one 30-nt overhang (**Fig. 6**). Complexes formed with these substrates remained stable for days after isolation by size-exclusion chromatography (SEC). Initial SAXS trials were performed with the full-length XPA and RPA proteins, but both had regions that retained a high degree of flexibility because they did not contact the other protein or the DNA substrate. Consequently, we prepared truncation constructs optimized for SAXS analysis: XPA_1-239_ and RPAΔ32NΔ70N (including deletion of the long RPA70N-RPA70A linker). High quality SAXS data were collected for complexes of XPA_1-239_ and RPAΔ32NΔ70N bound to both the 3’ and 5’ junction substrate. Analysis of these data (detailed description provided in Supplementary Material) revealed: (i) the solutions were free of aggregation; (ii) both complexes contain globular domains with a limited number of flexible loops and/or linkers; (iii) the ternary complex is stable and globular, which indicates that direct analysis of the distance distribution function, P(r), in terms of a molecular shape is feasible (**Table 3, Fig. S5**). *Ab initio* shape calculations were therefore performed with DENSS to obtain molecular envelopes for the two complexes (**Fig. 4, S6**).

**Table 3.** Summary of parameters obtained by SEC-SAXS for XPA-RPA-DNA complexes.

| **Constructs** | **D<sub>max</sub> (Å)** | **R<sub>g</sub> (Real) / R<sub>g</sub> (Reciprocal) (Å)** | **Porod Exponent** |
| --- | --- | --- | --- |
| XPA<sub>1-239</sub>/3' junction/RPAΔ32NΔ70N | 146.5 | 43.25/ 42.74 | 3.5 |
| XPA<sub>1-239</sub>/5' junction/RPAΔ32NΔ70N | 164.5 | 45.83/ 46.81 | 3.4 |
| XPA FL/3' junction/RPA FL | 196 | 51.8/ 49.75 | 3.0 |

**Figure 6:**
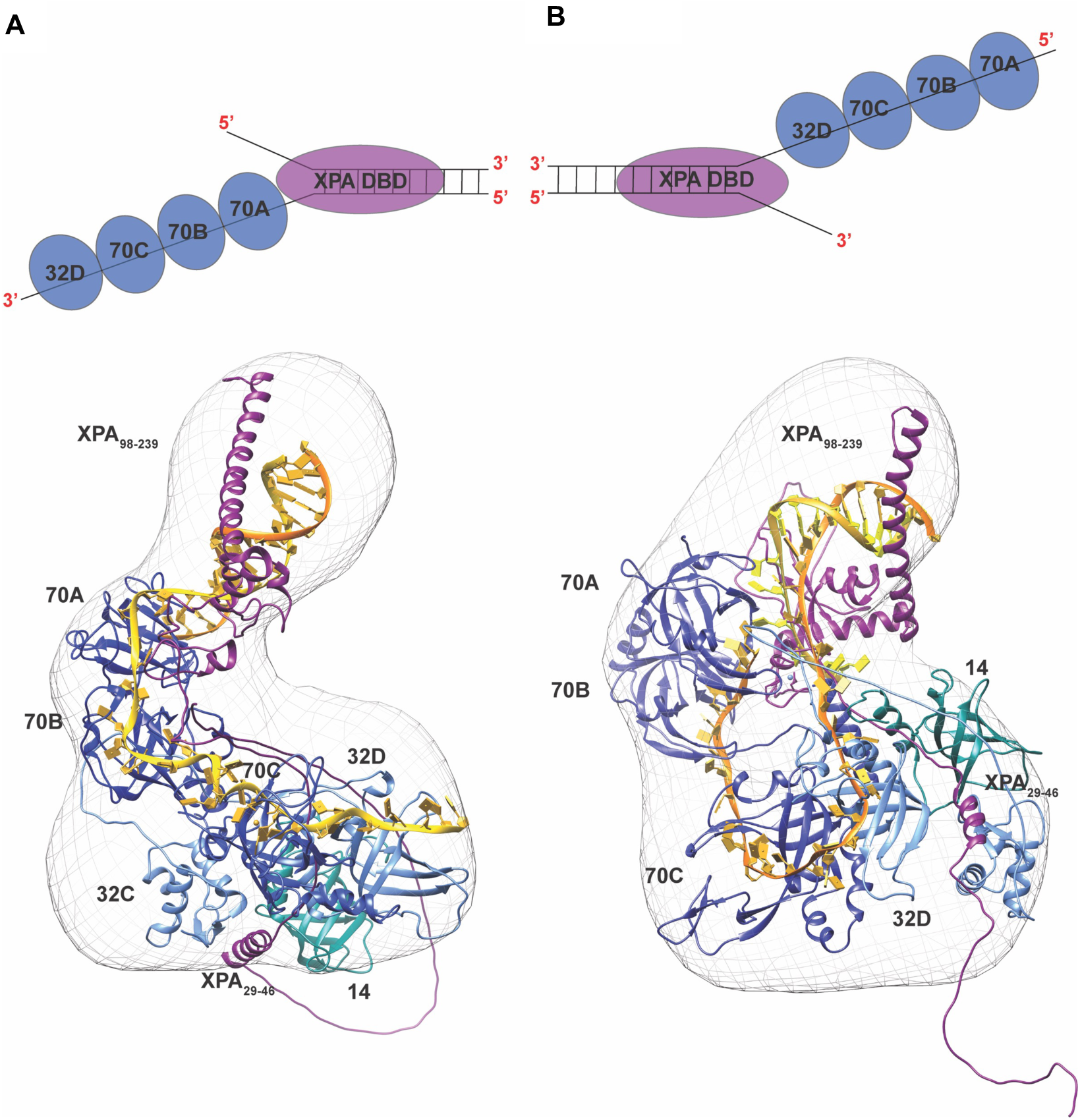
SAXS-based structural model of the complex of XPA_1-239_ and RPAΔ32NΔ70N complex engaged on two model NER substrates mimicking. **A**. 3’ss-ds junction and **B**. 5’ ss-ds junction. A schematic of XPA occupying the 3’ (A.) and 5’ (B.) DNA junctions and RPA subunits engaging the two DNA substrates is illustrated above the structures. Ribbon diagrams of the XPA–RPA bound to the model substrates with RPA subunits in shades of blue (RPA70ABC-dark blue, RPA32-light blue and RPA14-cyan), XPA DBD in purple and DNA in yellow for the 3’end and orange for 5’ end. The structures with the SAXS molecular envelopes place XPA DBD and RPA70AB interaction at the top portion of the envelope, revealing the reversal in the orientation of RPA32C/XPA_29-46_ interaction site for the two substrates. The final representative structural model is shown fit into the *ab initio* SAXS molecular envelope (mesh) generated by DENSS from the SAXS data.

A variety of previously determined XPA and RPA structures and SAXS molecular envelopes were used to generate structural models (see description in Supplementary Materials). The final representative models show reasonable fits to the corresponding experimental scattering profiles (3’ junction, χ= 2.8; 5’ junction, χ= 4.75) and the corresponding molecular envelopes. The model for the 3’ junction complex has an excellent fit to the data and results in a smoothly curved and compact arrangement of domains with the trimer core (RPA70C/32D/14) and RPA32C/XPA_29-46_ positioned within the larger lobe of the molecular envelope, and XPA-DBD/RPA70AB in the smaller extended lobe (**Fig. 6A**). The model for the 5’ junction complex has a similar curved arrangement of domains, but is a bit less compact (**Fig. 6B**). Importantly, both ss-dsDNA junction substrates are seen to be readily accommodated, consistent with our past observation that even with RPA bound to the undamaged strand there are no structural factors that inhibit XPA from binding to either junction of the NER bubble.

Despite their overall similarity, careful comparisons of the two models reveal some very clear differences, particularly in the orientation of the RPA domains. The positioning of the RPA32C-XPA_29-46_ contact is on opposite faces of the molecular envelope in the two complexes. Intriguingly, because RPA binds ssDNA in the 5’-3’ orientation, the path of the ssDNA in the complex with the 5’ ss-ds junction substrate is very different from that with the 3’ substrate. In the complex with the 3’ junction substrate, the ssDNA overhang mimicking the undamaged strand extends smoothly from the 3’ towards the 5’ junction. By contrast, the ssDNA in the complex with the 5’ junction substrate is highly bent and U-shaped to accommodate both the 5’-3’ polarity on the ssDNA and the RPA70AB-XPA interaction. As a result of this topological challenge, the ssDNA overhang mimicking the undamaged strand ends up remarkably close to the 3’ junction, placing the two ss/dsDNA junctions in close proximity. The implications of these observations for the structure of the NER pre-incision complex are discussed below.

## Discussion

In this study, we have analyzed the roles in NER of the two distinct interaction sites between XPA and RPA at the biochemical, structural, and cellular level. Our data show that both interactions are required for full NER activity and that mutations in the RPA-binding surfaces of XPA show additive defects, both physically and functionally. We propose that although similarly important for overall NER activity, the roles of these two contacts are distinctly different. Our data are consistent with a model in which the stronger XPA-RPA32C interaction is important to colocalize the two proteins, while the interaction between XPA-RPA70AB is needed for the proper positioning on the DNA bubble and licensing of the 5’ incision by ERCC1-XPF.

In our mechanistic models of how RPA functions, RPA32C and RPA70N serve as critical modules to recruit partner proteins to ssDNA bound by the multi-valent DNA binding apparatus (domains 70A, 70B, 70C, 32D), XPA_29-46_ was one the three homologous motifs first identified as characteristic of RPA32C binding sites in DNA repair proteins^20^. Subsequent studies have identified motifs with this signature sequence in a number of other RPA binding partners including SMARCAL1 and TIPIN ^48^. The highly modular RPA protein invariably engages its protein binding partners in a multi-valent mode with one interaction involving a primary recruitment module and a second weaker interaction that involves the tandem RPA70AB high affinity ssDNA binding domains. Despite having lower affinity, this interaction is critical to function because this is the interaction that positions the binding partner close to the DNA substrate. Hence, the multi-valent mode of binding to RPA fine-tunes XPA function; the interaction with RPA32C increases the local concentration thereby increasing the effective affinity of XPA-DBD for RPA70AB and stimulating their scaffolding function to position the NER nucleases XPF/ERCC1 and XPG. This multi-valent interaction mode provides an additional critical element to the NER machinery: the higher on-off rate associated with weak intrinsic affinity (e.g. between XPA-DBD and RPA70AB) is perfectly matched to the rapid structural transformations that are required for the progression through the NER trajectory ^14,49,50^. Our study furthermore informs ongoing efforts to examine RPA variant in human cancers^51^.

The functional data show that both contacts are required for NER function, although the mechanism for dysfunction when the one of the two contacts are defective is different. Our studies of the kinetics of association and dissociation of XPA at sites of UV damage show that XPA mutations in the RPA32C binding motif are defective in recruitment to the NER machinery, whereas mutations in the RPA70AB binding site, leading to increased residence times at UV damage, are defective in positioning of other NER factors and/or maintaining the stability of the pre-incision and incision complexes. The loss of recruitment via interaction with RPA32C is likely compensated by the fact that strong binding of RPA to the undamaged strand will not be affected, so free RPA can still find the strand via diffusion but its arrival on site will be slower. The defect in positioning via interaction with RPA70AB is likely compensated by interaction of XPA with other NER factors. In this case, XPA will not be held in place as effectively and so the NER machinery will be less efficient overall as the incision requires having all factors in place. We propose that these compensatory mechanisms are what provides the residual activity of the various XPA mutants observed in the functional assays.

One unanswered question is whether XPA and RPA are recruited simultaneously to the damage site. In prevailing general models, XPA and RPA are recruited together or XPA is recruited only after TFIIH creates the bubble and RPA is engaged. Reports of XPA, but not RPA, contacts with XPC and TFIIH supports a sequential model in which XPA is recruited first ^7,52^. RPA would then be recruited by the combination of its interaction with XPA and its strong affinity for the ssDNA generated as TFIIH unwinds the dsDNA around the lesion. However, cellular studies suggest that XPA and RPA can independently associate with sites of local UV damage ^53^. This leads to the question, whether RPA can only bind after the bubble is fully open, or whether RPA70AB could engage earlier as it binds tightly to as little as 8-nt of ssDNA? Interaction in this mode would fit well with a model in which RPA and XPA are recruited together to the bubble. Ongoing efforts to structurally examine the dynamic NER trajectory will reveal the intricate molecular mechanisms that govern the assembly and mechanics of the NER machinery.

We have previously shown that the XPA-DBD can interact with RPA70AB equally well bound to a model of the NER bubble with the ss-ds junction 3’ or 5’ to the lesion ^24^. This led us to question if XPA remains engaged to a specific ss-ds junction in the NER bubble or if it shifts from one to the other over the course of the NER trajectory. The structural analysis of the constructs containing both XPA-RPA contacts reported here revealed that the ability to engage either junction is retained. Nevertheless, studies of the isolated XPA-RPA complex are limited by the absence of the steric restrictions enforced by the other components of the NER complexes at various stages of the NER trajectory, the most significant of which is the 10-subumit TFIIH.

To determine if the binding of XPA and RPA at either junction would be compatible with constraints imposed by other NER factors, we generated models of XPA-RPA-TFIIH complex bound to DNA substrates with XPA located at either junction (**Fig. 7A, B**). Intriguingly, we find that both models can readily accommodate TFIIH without major structural reorganization. The close proximity of the two ss-ds junctions in both models is particularly noteworthy as it is consistent with curved nature of the RPA DNA binding apparatus and the potential for XPA to be bound at either junction at different points of the NER trajectory ^24^. As a further test of XPA binding at either junction, we used the XPA-RPA-TFIIH models to build the corresponding architectural models of the full NER pre-incision complex (**Fig. 7C**). These models are based on the following observations: (i) XPD binds the lesion-containing strand and lesion andXPB binds the duplex adjacent to the 5’ junction; (ii) in the cryo EM structure XPA interacts with TFIIH via XPD, XPB, as well as p8 and p52 ^19^; (iii) the RPA bound non-damaged strand has a U-shape ^34,54^; (iv) ERCC1-XPF is located at the 5’ junction, positioned for the 5’ incision ^55^; (v) XPG, by virtue of its interaction with TFIIH via p62 and XPD binds to the “backside” of TFIIH ^13,56^ and interacts with the duplex adjacent to the 3’ junction ^57,58^.

**Figure 7:**
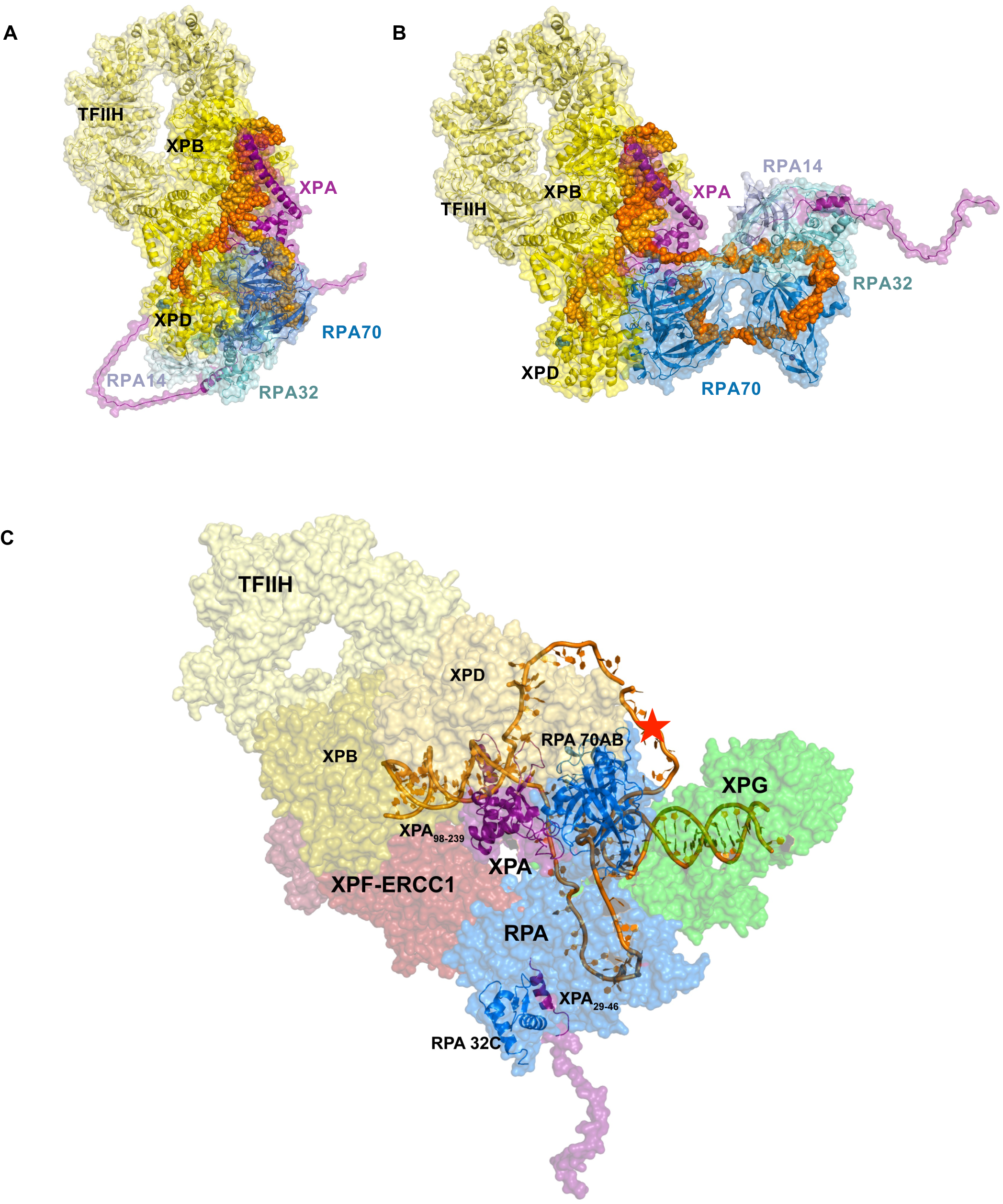
Structural models of NER complexes: **A./B**.The SAXS models of XPA-RPA-DNA are aligned by superimposing XPA DBD on the XPA-TFIIH cryo-EM structure (PDB ID:6R04). **A**. TFIIH-XPA-RPA bound to a 3’ ssDNA overhang junction. **B**. TFIIH-XPA-RPA bound to a 5’ ssDNA overhang junction **C**. Model of the pre-incision complex bound to the 5’ junction substrate. This model was prepared by aligning the cryo-EM structure of TFIIH-XPA (TFIIH in yellow) with our XPA-RPA-5’junction model (XPA in purple and RPA in blue) and placing the endonucleases XPF-ERCC1 (PDB: 6SXB) in red and XPG in green (PDB: 6Q0W) along the path of the dsDNA. The lesion is indicated as a star on the NER bubble (red). The two interaction points of XPA and RPA are highlighted as ribbons.

Current models of NER, based on the interaction of XPA with ERCC1-XPF to mediate 5’ incision and the cryo EM structure of the XPA-TFIIH complex ^19^with TFIIH at the 5’ junction, favor XPA at the 5’ junction. In the corresponding architectural model for the pre-incision complex, RPA70AB is bound on the undamaged strand close to the 3’ junction and the interaction between RPA70AB and XPA causes the 5 and 3’ junctions to be positioned close together. Perhaps the most striking feature of this model is the U shape of the two strands in the bubble. This arrangement is consistent with structures of the NER nucleases bending the DNA at the ss/dsDNA junction^55,57,58^. Our current work identifies the RPA70AB-XPA interactions as one of the key elements stabilizing this arrangement and shows that the details of the physical interaction between XPA and RPA are critically important to efficient and effective repair of lesions by the NER pathway.

## Acknowledgements

We thank Susan Tsutakawa (LBNL), Zachary Nagel (HSPH) and members of the Structural Biology of DNA Repair consortium for helpful comments and discussion. This work was supported by grants from the NCI (R01 CA218315 and P01CA092584 to ODS and WJC) and the Korean Institute for Basic Science (IBS-R022-A1 to ODS). MST instrumentation was acquired from National Institutes of Health through grant S10 OD021483. SAX studies were conducted at the Advanced Light Source (ALS), a national user facility operated by Lawrence Berkeley National Laboratory on behalf of the Department of Energy, Office of Basic Energy Sciences, through the Integrated Diffraction Analysis Technologies (IDAT) program, supported by DOE Office of Biological and Environmental Research. Additional support comes from the National Institute of Health project ALS-ENABLE (P30 GM124169). a High-End Instrumentation Grant S10 OD018483, the Structural Cell Biology of DNA Repair Machines program (P01 CA092584).

## Notes

### Competing Interest Statement

The authors have declared no competing interest.

### Summary of Updates

Correction of name spelling

