## Supplementary Materials for "Two Interaction Surfaces between XPA and RPA Organize the Preincision Complex in Nucleotide Excision Repair"

[1]Center for Genomic Integrity, Institute for Basic Science, Ulsan, Republic of Korea; Department of Biological Sciences,[2] Ulsan National Institute of Science and Technology, Ulsan, 44919, Republic of Korea. [3]Department of Biochemistry, [4]Center for Structural Biology, and [5]Department of Chemistry, Vanderbilt University, Nashville, TN 37232-7917, USA. [6]Department of Molecular and Cellular Oncology, The University of Texas MD Anderson Cancer Center, Houston, TX 77030, USA. [7]Biological and Systems Engineering Division, Lawrence Berkeley National Laboratory, Berkeley, CA.

**Table of Contents:**

**p. 2-5 Extended Materials and Methods Section**

**p. 5 Supplementary References**

**p. 6-12 Supplementary Figures**

#### EXTENDED METHODS

##### TFIIH purification

TFIIH core complex (XPB-PreScission-GFP, XPD, p62, p52, p44, p34, and p8) was cloned into MacroBac vector 438A [1]. The protein was expressed in Sf9 cells supplemented with 1 mM L-cysteine and 0.1 mM ferric ammonium citrate. Based on the XPB fusion with GFP, TFIIH was purified anaerobically using GFP-nanobody binder [2] covalently linked to agarose-beads (NHS agarose, Pierce/Thermo), eluted with PreScission protease, and purified using Superpose 6 (10/300) or Hi-Load Superdex 200 (16/600) for large-scale in 25 mM HEPES, pH 7.8, 150 mM NaCl, 50 mM KCl, 3% glycerol, and 3 mM 2-mercaptoethanol. The elution with TFIIH core complex (confirmed by SDS-PAGE gel and 420 nm absorption peak) was stored at – 80 °C until use.

##### SAXS data analysis

High quality SAXS data were collected for the complex of XPA and RPA with the 3' junction substrate. Analysis of the data revealed a monodisperse complex with a substantial degree of residual flexibility evident in the Kratky Plot and a value of 3.0 for the Porod exponent (**Table 3**). Importantly, the residual flexibility to this extent greatly limits structural interpretation of the data [3]. Both XPA and RPA have substantial flexible regions outside the two points of contact. For XPA this includes the C-terminal domain ($XPA_{240-273}$) and for RPA the disordered RPA32N domain ($RPA32_{1-42}$) and the globular RPA70N connected to RPA70A by a 68-residue flexible linker. We surmised that these regions are the origin of the residual flexibility observed in the SAXS data for the full-length proteins. Since these flexible regions do not contribute to the interactions between XPA and RPA or to binding the DNA substrate, we prepared the corresponding truncation constructs for further SAXS analysis: $XPA_{1-239}$ and RPAΔ32NΔ70N (including deletion of the long linker to RPA70N). As for the full-length proteins, mixing of truncated variants of XPA and RPA with the same DNA substrate provided a very stable complex.

SAXS data were collected first for the complex of $XPA_{1-239}$ and RPAΔ32NΔ70N bound to the 3' junction substrate. The SAXS profile and linearity of the Guinier plot (**Fig. S5B**) showed that the complex was free of aggregation in solution. The Kratky plot revealed both complexes contain globular domains and some flexible loops and/or linkers (**Fig. S5D**). Porod-Debye analysis, including the value of 3.5 for the Porod Exponent, indicated the ternary complex is stable and globular, and suggested that direct analysis of the distance distribution function, P(r), in terms of a molecular shape is feasible (**Table 3, Fig. S5F**). The primary peak centered at ~50 Å in the P(r) is attributed to distance distributions within the globular domains. The small feature in P(r) at ~120 Å is consistent with distances between regions of XPA and RPA that do not contact each other. SAXS data collected for $XPA_{1-239}$/RPAΔ32NΔ70N bound to the 5'junction substrate were of similar high quality, consistent with a well-formed and globular structure as reflected in the Porod Exponent of 3.4. The secondary shoulder in the P(r) is more prominent in the 5' junction complex at ~150 Å, suggesting there may be more flexibility within this complex. *Ab initio* shape calculations performed with DENSS provided molecular envelopes for the two complexes (**Fig. 6, S6**).

Previously determined SAXS analyses of the XPA-DBD/RPA70AB-DNA complex and of the RPA DNA binding apparatus (RPA-DBC) bound to ssDNA were used to interpret the molecular envelope for the 3'junction substrate complex (**Fig. S6**) [4,5]. The SAXS based structures of RPA-DBC bound to a 30-nt ssDNA oligomer and of XPA-DBD/RPA70AB bound to a Y-shaped ss-ds DNA junction were superposed by aligning the RPA70AB domains. Next, a homology model of the RPA32C bound to $XPA_{29-46}$ [6] was generated using Modeller [7]. The RPA32C/$XPA_{29-46}$ complex was then localized within the molecular envelope of the full complex using FoXSDock [4,8]. The representative structure selected for further analysis had the best combined score of $\chi$ fit to the scattering curve and energy. Disordered residues were added at the XPA N-terminus (1-28), between $XPA_{29-46}$ and XPA-DBD (47-97), and the linker residues between RPA32D and RPA32C ($RPA32_{172-199}$) using Modeller [9].The final representative structural model shows a reasonable fit to the experimental scattering profile ($\chi$= 2.8) and molecular envelope of the complex. This representative model exhibits a curved and compact arrangement of domains. The trimer core (RPA70C/32D/14) and the RPA32C/$XPA_{29-46}$ are positioned within the larger lobe of the molecular envelope, and XPA-DBD/RPA70AB in the smaller extended lobe (**Fig. 6A**).

The same overall approach was used interpreting the molecular envelope for the 5' junction substrate complex (**Fig. S6**). In this case, we were unable to achieve a good fit to the experimental data using the structures of the XPA-DBD/RPA70AB/DNA and RPA-DBC/DNA as was case for the 3' junction substrate complex. Rather, the best fit to data was achieved by docking the structures of the XPA-DBD/RPA70AB/DNA complex and the RPA trimer core (RPA70C/32D/14) with ssDNA bound extracted from SAXS-MD structure RPADBC bound to 20-nt ssDNA using FoXS Dock (32-33,43-44). The best representative scoring model was then further docked with the RPA32C/$XPA_{29-46}$ homology model. As for the 3'junction substrate complex, disordered residues were added using Modeller (45). This included at the N-terminus of XPA (1-28), between $XPA_{29-46}$ and XPA-DBD (47-97), the linker between RPA32D and RPA32C ($RPA32_{172-199}$), the linker between the RPA70B and RPA70C domains ($RPA70_{423-435}$), and the additional ssDNA (10 nucleotides) spanning from the 3' end of the ssDNA bound to RPA70B to the 5' end of the ssDNA bound to RPA 32C. The final representative structural model shows a reasonable fit to the experimental scattering profile ($\chi$= 4.75) and molecular envelope of the complex. This representative model exhibits a similar curved and compact arrangement of domains with the trimer core (RPA70C/32D/14) and RPA32C/$XPA_{29-46}$ positioned within the larger lobe of the molecular envelope, and XPA-DBD/RPA70AB in the smaller extended lobe (**Fig. 6B**).

### Figure S1.

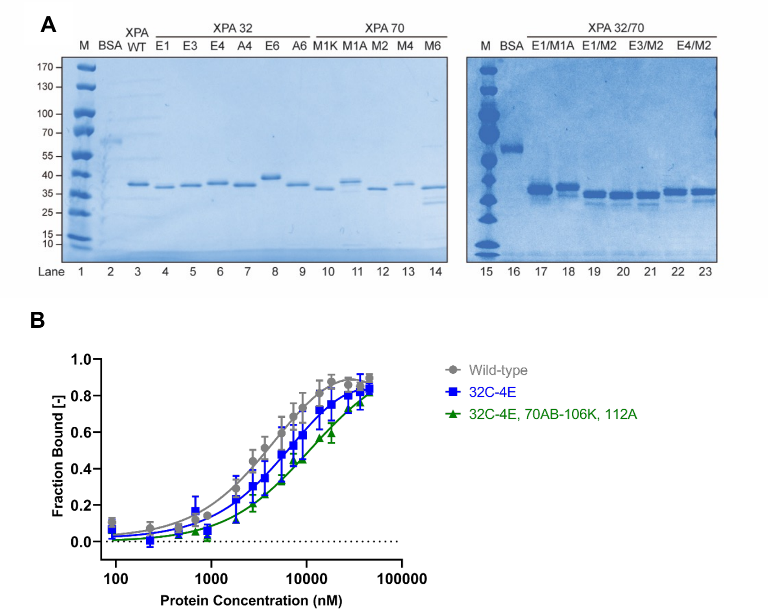


**Figure S1: DNA binding affinity of XPA wild-type, XPA-32-4E, and XPA-34E470-M2. A.** Purification of XPA-RPA mutant proteins. **B.** The mean and standard deviation of 3 independent MST measurements of $K_d$ values for the DNA substrate by wild-type, 32C-4E, and 32C-4E/70-M2 XPA were 5.4 ± 1.1 μM, 8.4 ± 2.3 μM, and 10 ± 3 μM, respectively. Error bars indicate the standard deviation ($N = 3$).

#### Figure S2.

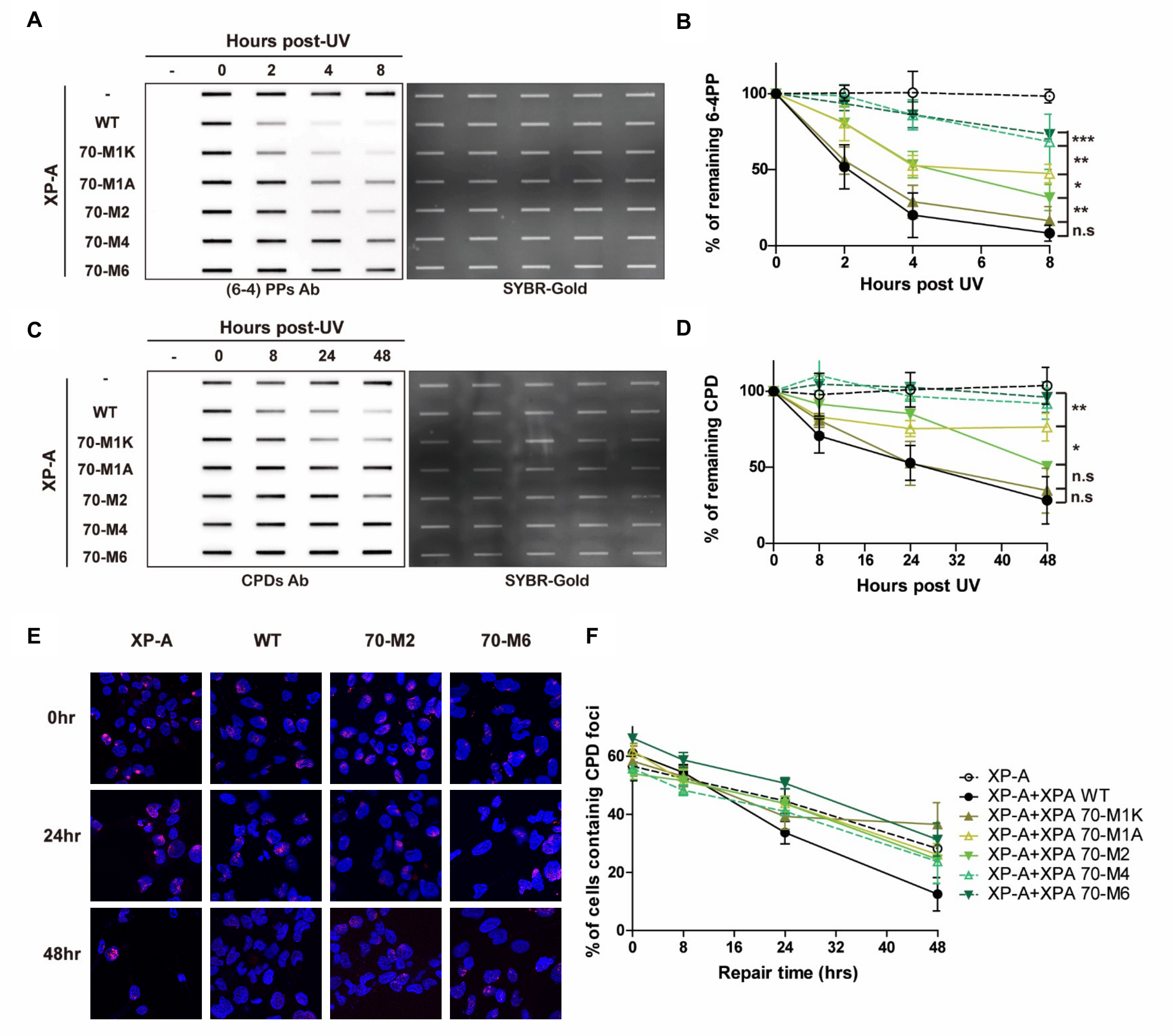


**Figure S2: (6-4) PP and CPD repair kinetics of XPA-RPA70 interaction mutants of XPA. A.** Determination of (6-4) PPs repair kinetics using slot-blot assays. Cells were irradiated with 5 J/m$^2$, genomic DNA isolated at indicated time point and the adduct levels determined using an anti-(6-4) PPs antibody. The right panel shows total DNA stained with EtBr. **B.** Quantification of A. Band intensity was normalized to the WT band at 0 hr. The data represent 3 independent experiments. The p-value was measured compared to XPA WT. **P*< 0.05, ***P*<0.01, ****P*<0.001. **C.** Determination of CPD repair kinetics. Slot blots were conducted as in A, except that an anti-CPD antibody was used. **D.** Quantification of C. Band intensity was normalized to the WT band at 0 hr and the data represent 3 independent experiments. The p-value was measured compared to XPA WT. **P*< 0.05, ***P*<0.01, ****P*<0.001. **E.** Representative figures of cells irradiated through a 5 μm micropore filter with UV irradiation 100J/m$^2$ and stained for CPD to measure repair kinetics. CPD foci are red and cell nuclei are stained blue with DAPI. **F.** Quantification of E. 100 cells were counted for each sample and the data represent at least 2 independent experiments.

#### Figure S3.

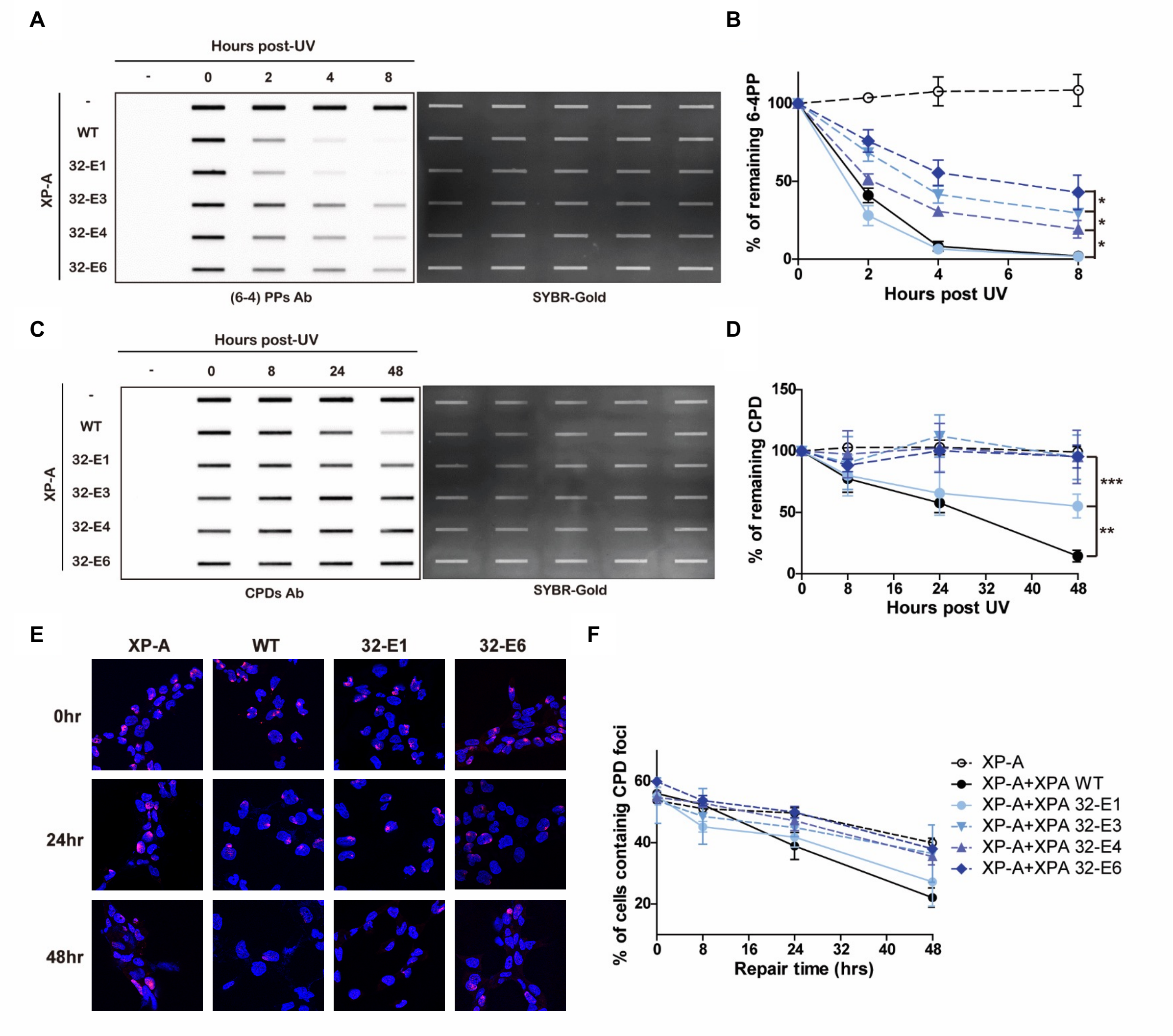


**Figure S3: (6-4) PPs and CPDs repair kinetics of RPA32 interaction mutants of XPA. A.** Determination of (6-4) PPs repair kinetics using slot-blot assays. Cells were irradiated with 5J/m$^2$ and genomic DNA isolated at indicated time point and the adduct levels determined using an anti-(6-4) PP antibody. The right panel shows total DNA stained with EtBr. **B.** Quantification of A. Band intensity was normalized to the WT band at 0 hr. The data represent 3 independent experiments. The p-value was measured compared to XPA WT. *$P$< 0.05, **$P$<0.01, ***$P$<0.001. **C.** Determination of CPD repair kinetics using slot-blot assays. Slot blots were conducted as in A, except that an anti-CPD antibody was used. **D.** Quantification of C. The data represent 3 independent experiments. The p-value was measured compared to XPA WT. *$P$< 0.05, **$P$<0.01, ***$P$<0.001. **E.** Representative figures of cells irradiated through a 5 µm micropore filter with UV (100J/m$^2$) and stained for CPD to measure repair kinetics. CPD foci are red and nuclei are stained blue with DAPI. **F.** Quantification of (E). 100 cells were counted for each sample and the data represent at least 2 independent experiments.

### Figure S4.

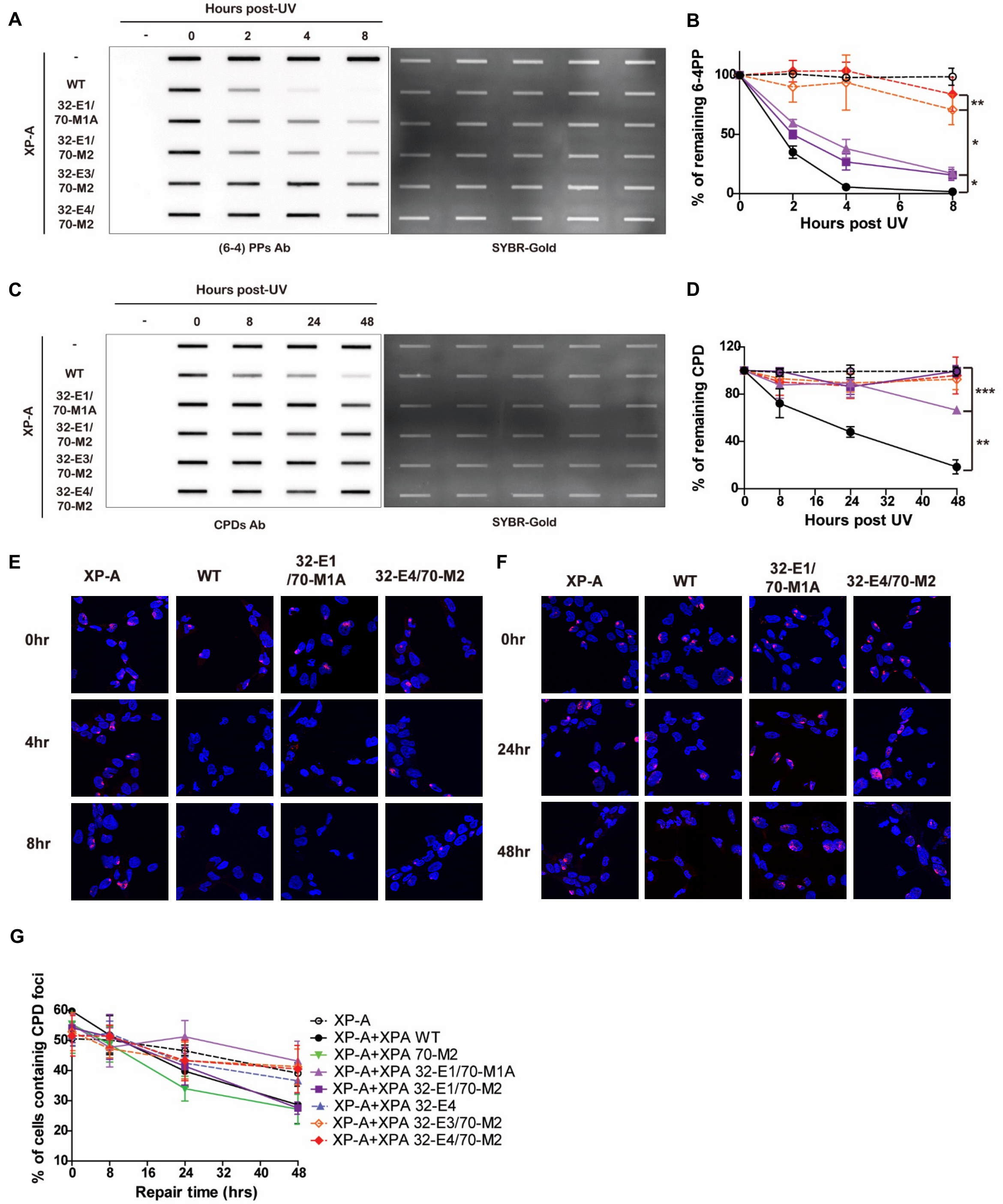
A
Hours post-UV
- 0 2 4 8
XP-A
-
WT
32-E1/ 70-M1A
32-E1/ 70-M2
32-E3/ 70-M2
32-E4/ 70-M2
(6-4) PPs Ab
SYBR-Gold
B
% of remaining 6-4PP
Hours post UV
C
Hours post-UV
- 0 8 24 48
CPDs Ab
SYBR-Gold
D
% of remaining CPD
Hours post UV
E
XP-A
WT
32-E1 /70-M1A
32-E4/70-M2
0hr
4hr
8hr
F
XP-A
WT
32-E1/ 70-M1A
32-E4/70-M2
0hr
24hr
48hr
G
% of cells containig CPD foci
Repair time (hrs)
XP-A
XP-A+XPA WT
XP-A+XPA 70-M2
XP-A+XPA 32-E1/70-M1A
XP-A+XPA 32-E1/70-M2
XP-A+XPA 32-E4
XP-A+XPA 32-E3/70-M2
XP-A+XPA 32-E4/70-M2

**Figure S4: (6-4) PPs and CPDs repair kinetics of RPA32/70 interaction mutants of XPA. A.** Determination of (6-4) PPs repair kinetics using slot-blot assays. Cells were irradiated with 5 $J/m^2$ and genomic DNA isolated at indicated time point and the adduct levels determined using an anti-(6-4) PP antibody. **B.** Quantification of A. Band intensity was normalized to the WT band at 0 hr and the data represent 3 independent experiments. The p-value was measured compared to XPA WT. *$P$< 0.05, **$P$<0.01, ***$P$<0.001. **C.** Determination of CPD repair kinetics using slot-blot assays. Slot blots were conducted as in A. except that an anti-CPD antibody was used. **D.** Quantification of C. The data represent 3 independent experiments. The p-value was measured compared to XPA WT. *$P$< 0.05, **$P$<0.01, ***$P$<0.001. **E.** Representative figures of cells irradiated through a 5 µm micropore filter with UV (100$J/m^2$) and stained for (6-4) PPs to measure repair kinetics. (6-4) PP foci are red and the cell nuclei are stained blue with DAPI. **F.** Representative figures of cells irradiated through a 5 µm micropore filter with UV (100$J/m^2$) and stained for CPD to measure repair kinetics. CPD foci are red and the cell nuclei are stained blue with DAPI. **G.** Quantification of F. 100 cells were counted for each sample and the data represent at least 2 independent experiments.

### Figure S5.

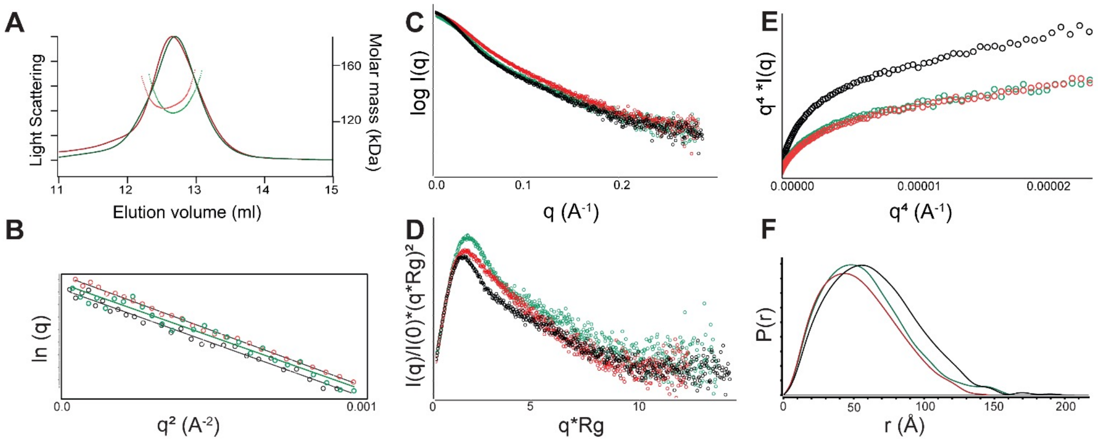


**Figure S5: SEC-SAXS analysis of XPA FL/3' junction/RPA FL complex (black), $XPA_{1-239}$/3'junction/RPAΔ32NΔ70N complex (red) and XPA1-239/5'junction/RPAΔ32NΔ70N complex (green)**. **A.** SEC-MALS trace showing the molar mass values (dots) observed across the elution profile. **B.** Guinier plots for the two complexes show that samples are free of aggregation. **C.** The full scattering profile for the three complexes. **D.** Kratky plots **E.** Porod plots. The Porod Exponent was estimated as 3.0 for the FL complex and 3.5 for the truncated complexes. **F.** Distance distribution analysis for the three complexes.

#### Figure S6.

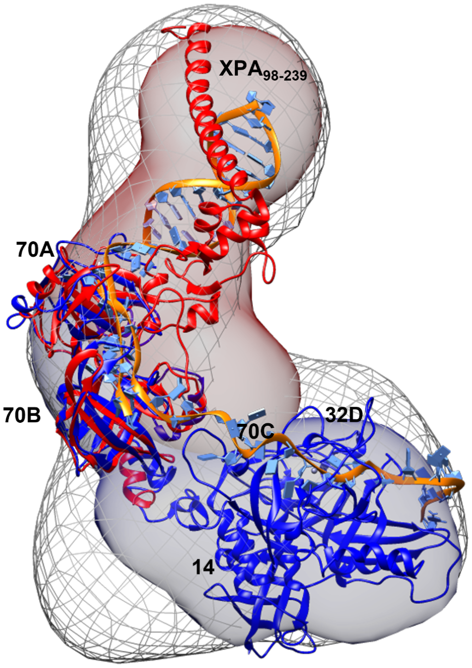


**Figure S6: Molecular envelope of $XPA_{1-239}$/3'junction/RPAΔ32NΔ70N complex (mesh) compared with SAXS molecular envelopes of RPA DNA binding core (blue) bound to 30nt ssDNA and $XPA_{98-239}$/DNA/RPAΔ32NΔ70N complex (red) prepared using DENSS.** The structures are superimposed at RP70AB domains. The unfilled region in the molecular envelope would correspond to RPA 32C/$XPA_{29-46}$ complex
